# Rad53 regulates the lifetime of Rdh54 at homologous recombination intermediates

**DOI:** 10.1101/2023.05.15.540757

**Authors:** Jingyi Hu, Bryan Ferlez, Jennifer Dau, J. Brooks Crickard

## Abstract

Rdh54 is a conserved DNA translocase that participates in homologous recombination (HR), DNA checkpoint adaptation, and chromosome segregation. *Saccharomyces cerevisiae* Rdh54 is a known target of the Mec1/Rad53 signaling axis, which globally protects genome integrity during DNA metabolism. While phosphorylation of DNA repair proteins by Mec1/Rad53 is critical for HR progression little is known about how specific post translational modifications alter HR reactions. Phosphorylation of Rdh54 is linked to protection of genomic integrity but the consequences of modification remain poorly understood. Here, we demonstrate that phosphorylation of the Rdh54 C-terminus by the effector kinase Rad53 regulates Rdh54 clustering activity as revealed by single molecule imaging. This stems from phosphorylation dependent and independent interactions between Rdh54 and Rad53. Genetic assays reveal that loss of phosphorylation leads to phenotypic changes resulting in loss-of-heterozygosity (LOH) outcomes. Our data highlight Rad53 as a key regulator of HR intermediates through activation and attenuation of Rdh54 motor function.

## Introduction

DNA double strand breaks (DSBs) are a dangerous type of DNA lesion that if unrepaired result in cell death or loss of genomic integrity (1). DSBs result from normal DNA metabolism, and from exposure to mutagens within the environment (1–3). This makes double strand break repair (DSBR) critical for the maintenance of genomic integrity. DSBR is a multi-tier decision making process in which different pathways can be used to repair the break depending on the phase of the cell cycle. The two primary DSBR pathways are non-homologous end joining (NHEJ) and homologous recombination (HR) (1). HR is a template based DSBR pathway that is primarily active during S/G2 phase of the cell cycle and meiosis (2,4–7). HR outcomes are controlled by the selection of a donor DNA template (8–13), and the alignment of both sides of the DSB. Known as DNA second end capture (14–18).

In eukaryotes template selection is coordinated by the recombinase Rad51 (7,19,20), which catalyzes the alignment of ssDNA (recipient DNA) from upstream of the DSB to a matching dsDNA (donor DNA) elsewhere in the genome in a process called the homology search. In *Saccharomyces cerevisiae*, Rad51 is aided in alignment of recipient and donor DNA by several accessory factors including Rad52, Rad54, Rdh54, and Rad55/57 (21–31). In mitotic HR, the sister chromatid is the primary template used for repair and results in the highest fidelity outcome (32,33). However, in rare cases the homologous chromosome is used as the template for repair. When the homologous chromosome is used as the template there is potential for gene conversion (34), which can result in a loss-of-heterozygosity (LOH) (35–38) (**Figure 1A**). LOH is a driver in many human cancers making understanding mechanisms that regulate these outcomes important.

**Figure 1:**
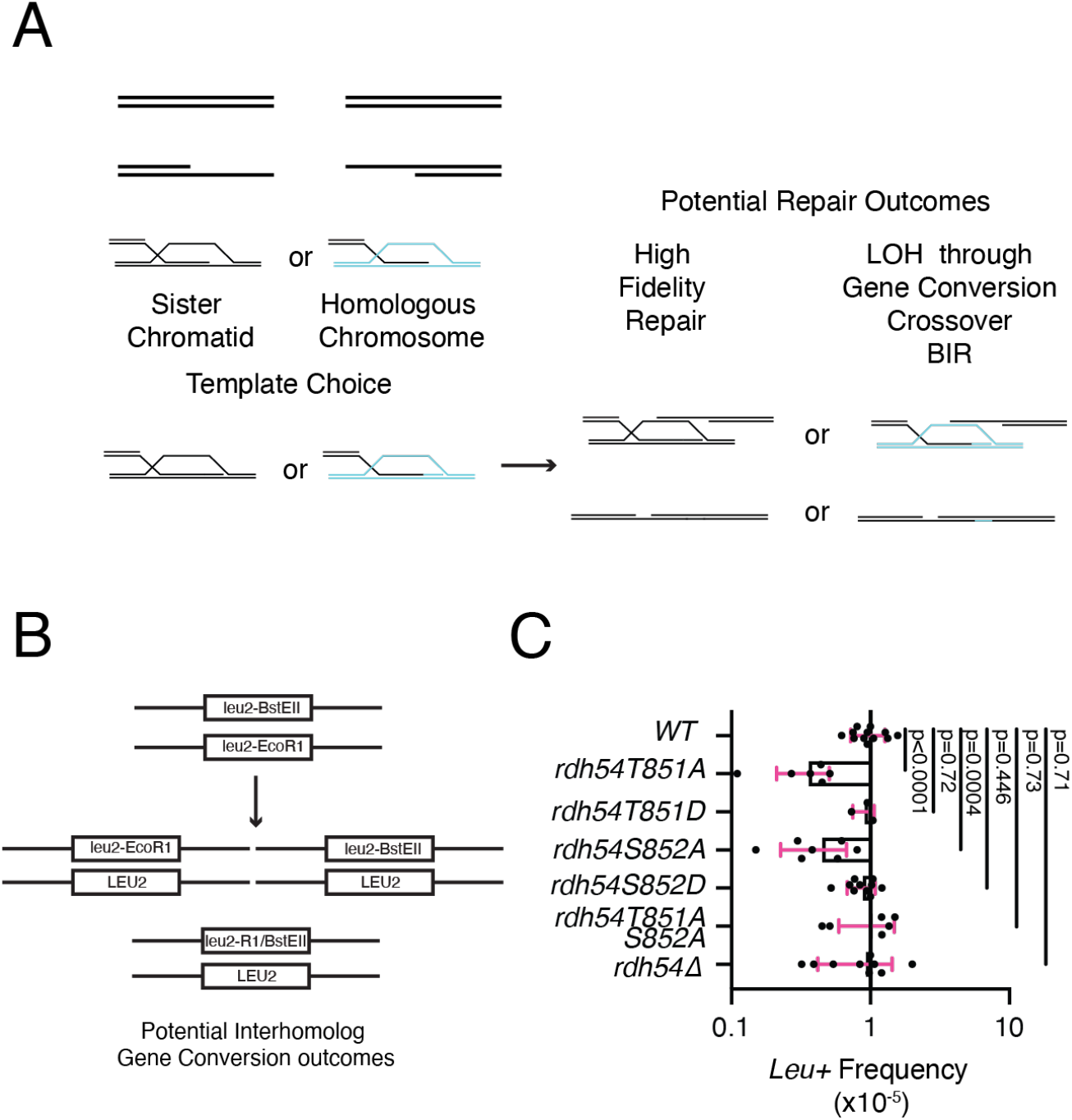
Rdh54 mutants’ effect on interhomolog recombination. **(A).** Schematic diagram illustrating how chromosome template selection can influence the outcome of HR leading to LOH outcomes. **(B).** Assay designed to identify gene conversion between homologous chromosomes. The assay is dependent on the formation of an *LEU2* from two inactive *leu2* alleles. **(C).** Comparison of interhomolog gene conversion frequencies for WT, *rdh54T851A*, *rdh54T851D*, *rdh54S852A*, *rdh54S852D*, *rdh54T851AS852A,* and *rdh54Δ*. The crossbar represents the mean, and the error bars represent the standard deviation of at least three independent experiments from multiple crosses. Statistical analysis was performed using a non-parametric Mann-Whitney test.

LOH outcomes result from exchange of genetic information between homologous chromosomes. These outcomes are dictated by competing pathways that depend on resolution of intermediates that form during HR mediated repair. These pathways include synthesis-dependent strand annealing (SDSA) (34), the classical double Holliday junction pathway (dHJ) (4,39), and break-induced replication (BIR) (17,40–44). Of these pathways the dHJ and BIR have the potential to create large scale rearrangements of chromosomes through reciprocal and non-reciprocal exchange of DNA, respectively (5,44,45). Outcomes during HR are strongly influenced by the timing of each step, which is under the regulation of DNA damage kinases (46). These kinases also play established roles in slowing cell cycle progression. As a result, their activation provides more time for the execution of DNA repair before cell division and reduces the passage of damaged DNA (46). In *S. cerevisiae*, the sensor kinase Mec1 plays a predominant role in activation of damage signaling kinases (47), with a major downstream target being the effector kinase Rad53 (47–50). Rad53 phosphorylates proteins within the HR pathway (46,51), at the replication fork (52–54), and at the kinetochore where it regulates chromosome division (55–58). These kinases are believed to influence DNA repair pathway choice during DSBR.

Rdh54 is a target of the Mec1/Rad53 signaling axis (59), but it is unknown how phosphorylation of Rdh54 regulates function. Rdh54 is a DNA motor protein that uses the power of ATP hydrolysis to physically move along dsDNA and is directly recruited to HR intermediates through interactions with Rad51 (28,30,60,61). Rdh54 has been shown to stabilize Rad51 bound to ssDNA and displacement loops, and to disrupt Rad51 bound to dsDNA (62–65). Rdh54 forms a homo-oligomer through self-association facilitated by a flexible disordered N-terminal region (61,66,67). However, it remains unclear if this behavior has biological function. Rdh54 has been genetically implicated in regulating CO/NCO outcomes (26,68), BIR outcomes (62), interhomolog gene conversion (68,69), and template switching associated with BIR (70,71). Importantly, Rdh54 is not required for completion of these pathways, and instead appears to only influence HR choices indirectly. It has also been suggested that Rdh54 activity is context dependent.

Here we have identified and characterized Rdh54 mutants that affect communications between monomers of Rdh54. Mutations in this region lead to changes in Rdh54 clustering behavior on dsDNA, translocation activity on dsDNA, and interactions with the effector kinase Rad53. We also find this region to be a target of Rad53 whose kinase activity appears to down regulate the translocation activity of Rdh54 on dsDNA. Importantly, mutations in this region have phenotypic consequences on the distribution of LOH outcomes associated with HR. From this data we have classified Rdh54 mutations as translocation fast and slow versions of the enzyme linking these classes of mutants to different phenotypic outcomes. Finally, we have developed a model where Rdh54 activity at DNA repair sites is controlled by a dynamic equilibrium that balances recruitment of Rdh54 by Rad51 and dissociation of Rdh54 through translocation along dsDNA. This dynamic equilibrium helps regulate HR outcomes by a kinetic mechanism which is coordinated by Rad53 activity.

## Materials and Methods

### Yeast strain construction

Most of the *S. cerevisiae* strains used in this study were W303. All recombination outcome experiments were performed in the W303 background. Strains for spot assay experiments were BY4741 and were generated by transformation of *rdh54*Δ strains with pRS415 plasmids. The genotypes for all strains used in this study can be found in **Supplemental Table 1**. Plasmids for generation of strains can be found in **Supplemental Table 2**.

### Yeast spot growth assay and colony growth

For complementation spot assays, overnight cultures were diluted back to an OD_600_ of 0.2 and then allowed to grow to an OD_600_ of 1.0. Cells were then serially diluted and manually spotted on Synthetic Complete (SC) -Leu plates containing either no drug, 0.01% MMS or 25 µM CPT. Plates were incubated at 30°C for 2-3 days and imaged at 24, 48, and 72 hours. For colony growth, overnight cultures were diluted 100-fold and then allowed to grow 2 hours. Then 0.01% MMS or an equal volume of sterilized H_2_O was added to the cultures and allowed to grow 4 hours. Colony forming units (CFU) were determined by plating 1000-fold diluted cultures on YPD + adenine sulfate plates. The plates were scored for colony growth and survival percentage was calculated by dividing the colonies from MMS-treated culture by the untreated culture.

### Interhomolog gene conversion assay

Yeast strains of JBC206 and JBC207 and derivatives were crossed (**Supplemental Table 1**), and the diploids were selected on YNB (-Ura/-Trp) + 2% dextrose and allowed to grow for 2-3 days. Fresh diploid colonies were then grown for 15 hours in YPD. 30 µl of saturated culture was plated on YNB (-Leu) + 2% dextrose and 30 µl of culture diluted 1:10,000 was plated on YNB (SC) + 2% dextrose and allowed to grow for three days. The colonies on each plate were then counted. The data was analyzed by dividing the adjusted number of colonies on the -Leu plate by the colonies on the SC plate. This generated a frequency. The mean and standard deviation of the data were determined using multiple different colonies from multiple different crosses. Statistical comparisons were also performed using a non-parametric Mann-Whitney test for significance. This did not yield different results from the standard t-test.

### Red/White recombination assay

The WT strain used in this assay as well as the procedure for diploid formation are described here (45,62). The genotypes for modifications to these strains can be found in **Supplemental Table 1**. The assay was performed by growing the appropriate strain overnight in YP + 2% raffinose. The next day cells were diluted to an OD_600_ of 0.2 and allowed to reach an OD_600_ of 0.4-0.5 followed by the expression of *I-Sce1* through the addition of 2% galactose. Cells were allowed to grow for an additional 1.5 hours, after which they were plated on YPD and allowed to grow for 48 hours. After 48 hours they were placed at 4 °C overnight to allow further development of red color. The number of white, red, and sectored colonies was then counted followed by replica plating onto YPD + hygromycin B (200 µg/ml) and YPD + nourseothricin (67 µg/ml, clonNat) for analysis of recombination outcomes. Strains were also replica plated on YNB (-Ura/-Met) + 2% dextrose to insure proper chromosome segregation (**Supplemental Table 4**). The data was analyzed by counting colonies that were sectored, and then counting colony survival on different antibiotic sensitivities. The data for each category was then divided by the total population of sectored colonies. The standard deviation between biological replicates analyzed for at least three independent experiments from different crosses.

### Template switching assay

The WT strains used for this study were described in (70). The genotypes for modification of these strains can be found in **Supplemental Table 1**. The assay was performed by growing the appropriate strain in YP + 2% raffinose overnight. The next day cells were diluted in YP + 2% raffinose to an OD_600_ of 0.2 and allowed to grow for four hours. The cells were then treated with 2% galactose for 3 hours and plated on YNB (-Ura) + 2% dextrose. CFU were determined by plating a dilution of the culture on non-selective media. After three days the plates were scored for colony growth. The Ura*+* frequency was determined by dividing the adjusted CFU from the selective plate by the number of colonies on the non-selective plate. The mean and standard deviation were calculated for three independent experiments.

### Co-immunoprecipitation experiments

A starter culture was grown in 5 mL YPD for 10 hours. The culture was then inoculated into 100 mL of YP + 3% glycerol + 2% sodium lactate and allowed to grow overnight until the OD_600_ was ∼1.0. Then 2% galactose (final concentration) was added to the culture to induce DSBs. In an uninduced control group, 2% dextrose was added. The cells were allowed to grow for 2 hours and harvested by centrifugation. Cell pellets were washed twice with 1xPBS (137 mM NaCl, 2.7 mM NaCl, 1.8 mM KH_2_PO_4_, 10 mM Na_2_HPO_4_) + 0.2 mM PMSF, and frozen at #x2212;80 °C for at least 30 minutes. Cells were then thawed and resuspended in lysis buffer (20 mM Tris-Cl pH 7.5, 300 mM NaCl, 0.01% NP-40, 10% glycerol,0.2 mM PMSF, Protease inhibitor cocktail). Glass disruptor beads were added, and the cells were bead beat with 1 minute on and 1 minute off for a total of 8 cycles. The cellular debris was then spun down at ∼1,100 x g for 10 minutes at 4 °C. The supernatant was then centrifuged at 26,712 x g for 10 minutes at 4 °C. The supernatant was collected as whole cell extract and incubated with anti-RAD53 (Abcam, ab104232) pre-coated Dynabeads Protein G (Invitrogen, Cat No. 10003D) overnight at 4 °C. To prepare the coated beads, they were first washed with PBST (PBS + 0.1% Tween-20) 3 times, then resuspended with PBST and antibody and incubated at 4 °C for 2 hours. After incubation, the antibody was removed, and the beads were washed with PBST 3 times. Then the beads were ready to be used. For each 1 mL of cell lysate, 1 µL of anti-RAD53 antibody and 15 µL of Dynabeads were used. The immunocomplexes were washed with wash buffer (20 mM Tris-Cl pH 7.5, 500 mM NaCl, 0.01% NP-40, 10% glycerol) 3 times and transferred to a new tube. The whole cell extract and immunocomplex were dissolved in SDS loading buffer. The samples were loaded on an SDS-PAGE, and the proteins were transferred onto a nitrocellulose membrane. The membrane was stained with Ponceau S staining solution (5% glacial acetic, 0.1% Ponceau S) and blocked with 5% non-fat dry milk in TBST (2.4 g Tris, 8.7 g NaCl, 1 mL Tween-20 in 1 L H_2_O, pH 7.5) for 1 hour. The membrane was then incubated with the primary antibody (anti-HA, Roche, 11583816001, 1:1000) in TBST + 5% non-fat dry milk overnight at 4 °C followed by incubation with secondary antibodies (Goat anti-Mouse IgG, Invitrogen, Cat No. A16072, 1:3000) at room temperature for 1 hour. The immunoblots were visualized with enhanced chemiluminescence.

### Protein expression and purification

6xHis-Rdh54, 6xHis-Rdh54-mCherry, and Rdh54 mutants were purified as follows. Rosetta 2 Bl21 (DE3) cells bearing pET15-6xHis-Rdh54 expression plasmids were grown to an OD_600_ between 0.6 and 0.8 at 37 °C. The temperature was lowered to 16 °C and cells were induced with 0.5 mM IPTG overnight. Cells were harvested by centrifugation and stored at −80 °C. Cells were resuspended in buffer A (25 mM Tris-HCl pH 7.5, 1000 mM NaCl, 10 mM Imidazole, 0.01% NP-40, 10% Glycerol, 5 mM 2-Mercaptoethanol, 0.2 mM PMSF, and Protease Inhibitor cocktail) and lysed by freeze thaw. Cell lysate was sonicated on ice for 10 cycles with 65% duty cycle and 15 sec on and 45 sec off. Lysate was centrifuged at 10,000 x g for 45 minutes at 4 °C. Clarified extract was then mixed with pre-equilibrated His-Pur Nickel-NTA resin and incubated in batch for 1 hour at 4 °C. After 1 hour the resin was centrifuged and washed 2x with buffer A, followed by 2x washes with buffer B (25 mM Tris-HCl pH 7.5, 200 mM NaCl, 10 mM Imidazole, 0.01% NP-40, 10% Glycerol, 5 mM 2-Mercaptoethanol, 0.2 mM PMSF, and Protease Inhibitor cocktail). The resin was then added to a disposable column, and the protein was eluted in buffer C (25 mM Tris-HCl pH 7.5, 200 mM NaCl, 200 mM Imidazole, 0.01% NP-40, 10% Glycerol, 5 mM 2-Mercaptoethanol, 0.2 mM PMSF, and Protease Inhibitor cocktail). Fractions were analyzed by SDS-PAGE and the peak fractions were pooled. The protein was further purified using a strong cation exchange column (HiScreen SP FF). The column was resolved using a 20-100% gradient of buffer D (25 mM Tris-Cl pH 7.5, 0 mM KCl, 1 mM EDTA, 0.01% NP-40, 5 mM 2-Mercaptoethanol, 10% Glycerol) and buffer E (25 mM Tris-Cl pH 7.5, 1000 mM KCl, 1 mM EDTA, 0.01% NP-40, 5 mM 2-Mercaptoethanol, 10% Glycerol). Fractions were analyzed by SDS-PAGE. Peak fractions were pooled and concentrated by centrifugation at ∼1,100 x g at 4 °C using a 10 kDa MWCO Vivaspin 6 column. Protein was finally purified on a 120 mL S300 column (HiPrep Sephacryl S-300 HR) in SEC buffer (30 mM Na-Hepes pH 7.5, 400 mM NaCl, 1 mM EDTA, 0.01% NP-40, 5 mM 2-Mercaptoethanol, 10% Glycerol). Peak fractions were pooled, and the protein was concentrated as described above. Purified protein was stored at −80 °C. 6xHis-Rad53, 6xHis-GFP-Rad53, 6xHis-Rad53D339A, and 6xHis-GFP-Rad53D339A were purified as previously described (52). Plasmids for protein expression can be found in **Supplemental Table 2**.

### In vitro kinase reactions

In vitro kinase assays were performed in Rad53 kinase buffer (25 mM Na-Hepes pH 7.5,150 mM NaOAc, 10% Glycerol, 10 mM MgCl_2_, 10 mM ATP, 0.2 mM Na Orthovanadate). 1 µM Rdh54 was incubated with 0.1 µM Rad53 for the defined periods of time. Reactions were quenched with SDS-PAGE loading buffer and resolved on a 10% SDS-PAGE impregnated with phostag reagent and 10 mM MnCl_2_. The gel was stained with Coomassie Brilliant blue and the shift in Rdh54 was directly observed. Mass spectrometry analysis was performed on Rdh54 samples phosphorylated *in vitro* that were excised from a Coomassie-stained SDS-PAGE.

### ATPase assay

To measure ATP hydrolysis rates, a commercially available ADP-GLO kit was used. ATP hydrolysis reaction were performed in HR buffer (20 mM Tris-OAc, 50 mM NaCl, 10 mM MgOAc_2_, 200 ng/µl BSA, 1 mM DTT and 10% Glycerol) and contained 1 mg/ml sheared salmon sperm DNA, and 100 nM Rdh54.

### Flow cell construction

To generate flow cells, metallic chrome patterns were deposited on quartz microscope slides with pre-drilled holes for microfluidic line attachment by electron beam lithography (72). After metal deposition microscope slides were converted to flow cells by adhering a piece of double-sided tape to the side of the microscope slide with the metal barriers. A channel was created by covering the two-sided tape with a small piece of paper in between the two drill holes. The paper was excised to create the flow chamber and a glass coverslip was fixed to the tape. The chamber was sealed by heating to 165°C in a vacuum oven at 25 mm Hg for 60 minutes. Flow cells were then completed by using hot glue to fit IDEX nano ports over the drill holes on the opposite side of the microscope slide from the coverslip.

### Single molecule experiments

All single molecule experiments were conducted on a custom-built prism-based total internal reflection microscope (Nikon) equipped with a 488-nm laser (Coherent Sapphire, 100 mW), a 561-nm laser (Coherent Sapphire, 100 mW), and two Andor iXon EMCCD cameras. DNA substrates for DNA curtains experiments were made by attaching a biotinylated oligo to one end of the 50 kb Lambda phage genome, and an oligo with a digoxygenin moiety on the other. This allowed double tethering of the DNA between chrome barriers and chrome pedestals, as previously described (72,73). Flow cells were attached to a microfluidic system and sample delivery was controlled using a syringe pump (KD Scientific). Two color imaging was achieved by two XION 512×512 back-thinned Andor EM-CCD cameras and alternative illumination using a 488-nm laser and a 561 nM laser at 25% power output. The lasers were shuttered resulting in a 200 msec delay between each frame. Images were collected with a 200 msec integration time. Rdh54-mCherry translocation velocity and distances were measured in HR Buffer (20 mM Tris-OAc, 50 mM NaCl, 10 mM MgOAc_2_, 200 ng/µl BSA, 1 mM DTT).

### Analysis of dsDNA translocation

The velocity and track length for Rdh54-mCherry molecules were measured by importing raw TIFF images as image stacks into ImageJ. Kymographs were generated by defining a 1-pixel wide region of interest (ROI) along the long axis of individual dsDNA molecules. Data analysis was performed from the kymographs. The start of translocation was defined when the Rdh54-mCherry molecule moved > 2 pixels. Pauses were defined as momentary stalls in translocation that lasted 2-4 frames. Termination was defined by molecules that did not move for >10 frames.. Velocities were calculated using the following formula [(|Y_f_-Y_i_|)*1,000 bp/[|X_f_-X_i_|])*frame rate]; where Y_i_ and Y_f_ correspond to the initial and final pixel position and X_i_ and X_f_ correspond to the start and stop time (in seconds). Graphs of individual velocity and distances travelled were plotted in GraphPad Prism 9. The mean was determined from these graphs and significant differences between mutants was determined by performing a student’s t-test of this data.

### Photobleaching measurements and analysis of Rdh54 cluster size

Photobleaching measurements were made by binding Rdh54-mCherry to the dsDNA and then observing the reaction without shuttering of the 561 nM laser at 25% power output (25 mW). Images were collected with 200 msec integration time. Kymographs were generated from individual DNA molecules and were visually inspected for irreversible changes in signal intensity. This was quantified by measuring the signal intensity over time and identifying defined drops. These drops in intensity were used to create a distribution of photobleaching steps. The intensity of individual drops was used to create a distribution to evaluate the mean intensity drop which correlates with the intensity for an individual fluorophore. Cluster size was then determined by subtracting the global background from the flow cell from each cluster, and then dividing by the mean intensity drop for an individual fluorophore. This created an estimated number of molecules per foci. Populations of bound Rdh54 under different conditions were statistically compared for significance using a non-parametric Mann-Whitney test in Graph Pad Prism 9.

### Rad53 binding experiments

For Rdh54 binding experiments, 1 nM of 6xHis-Rdh54-mCherry was mixed with 6xHis-GFP-Rad53 and injected onto pre-formed DNA curtains. Initial binding was monitored with a frame rate of 1 frame/3 sec. Data were then processed as described above. The number of Rdh54 molecule bound by GFP-Rad53 were then scored for binding. For functional assays Rdh54-mCherry, Rdh54T851A-mCherry, or Rdh54S852A-mCherry were treated with catalytic amounts of Rad53 for 60 minutes in HR buffer. Translocation was then monitored by single molecule imaging and analyzed as described above.

## Results

### A Rdh54 phosphorylation mutant displays enhanced sensitivity to DNA damage

Rdh54 is a known target of the Mec1/Rad53 kinases (59). However, the effects of Rdh54 phosphorylation during DNA repair are unknown. To identify potential phosphorylation sites, we used the SuperPhos database (74). Nine phosphorylation sites were identified in Rdh54: S19, S51, S165, S386, S619, S790, S791, T851, and S852 (**Supplemental Figure 1A**). These numbers are based on Rdh54 from the W303 genetic background. In the reference strain (S288C) there is an additional thirty-four amino acids in the N-terminal region of Rdh54. To map these sites to the reference strain, add thirty-four to the amino acid number used in the W303 strain. For example, S19 in W303 is S53 in the reference strain. Of these nine sites, one was part of a Rad53 consensus sequence (S852) (75), and three were high confidence sites. High confidence sites are defined as >10 peptides identified in the proteomic analysis (S51, S619, and S852) (**Supplemental Figure 1A**). We initially converted all suspected phosphorylation sites to alanine and tested them for their ability to complement methyl-methanosulfonate (MMS) and camptothecin (CPT) sensitivity phenotypes in *rdh54Δ* haploid yeast strains. In addition to the serine/threonine to alanine mutations, we also tested the *rdh54K318R* allele which harbors a mutation in the Walker A box of Rdh54 preventing hydrolysis of ATP. We used this mutant because it has damage sensitivity phenotypes that are worse than the *rdh54Δ* strain (30). Most of the S to A mutations complemented the weak MMS and CPT phenotypes associated with the *rdh54Δ* (**Supplemental Figure 1B**). However, one mutant, *rdh54T851A*, failed to complement sensitivity to CPT and MMS (**Supplemental Figure 1B**).

To validate the screen, we integrated *rdh54T851A* into the genome by gene replacement and tested for MMS sensitivity. Because we were testing the potential impact of phosphorylation we also included *rdh54T851D, rdh54S852A, rdh54S852D*, and *rdh54T851A,S852A*. The S852 residue was included because it is the most highly phosphorylated site on the SuperPhos database and is also a part of a Rad53 consensus sequence. We measured sensitivity by scoring percent survival as compared to WT. Under the conditions used in our experiments WT strains had a survival percentage of 41.6+/-7 (**Supplemental Figure 2A**). In contrast several Rdh54 mutants showed small but reproducible defects in survival. For the *rdh54T851A*, *rdh54T851D*, and *rdh54S852D* strains the survival percentages dropped to 25+/-6, 25+/-4, and 24+/-5.6, respectively (**Supplemental Figure 2A**). These values were comparable to the survival percentage of the rdh54Δ strains (26+/-2%). In contrast, the *rdh54S852A* and *rdh54T851A,S852A* alleles were not significantly different from the WT with 40+/-4 and 48+/-8 percent survival respectively (**Supplemental Figure 2A**). From this data we conclude that mutations in this region of Rdh54 can cause weak MMS sensitivities. We additionally validated that these mutants were expressed at comparable levels in cells by western blot (**Supplemental Figure 2B**).

### Mutation of phosphorylation sites alters interhomolog recombination

Genetic experiments have implicated *RDH54* in regulating interhomolog (IH) gene conversion, and LOH outcomes (68,69). Therefore, we tested *rdh54T851A*, *rdh54T851D, rdh54S852A*, *rdh54S852D,* and *rdh54T851A,S852A* alleles for their ability to promote LOH gene conversion. We used an assay that measures the gene conversion frequency of two inactive *leu2* alleles (*leu2-EcoRI* and *leu2-BstEII*) located on homologous chromosomes into an active *LEU2* gene (**Figure 1B**). The recombination outcomes that cause conversion to Leu+ include the generation of a *LEU2/leu2-EcoRI* gene pair, the generation of a *LEU2/leu2-BstEII* gene pair, or the generation of a *LEU2*/*leu2-EcoRI-BstEII* gene pair (**Figure 1B**). This last outcome results from reciprocal exchange of DNA between chromosomes. It should be noted that there are other possible outcomes that can occur due to recombination between heteroalleles. These include non-reciprocal exchange of DNA that does not result in the formation of a *LEU2* allele and instead results in a *leu2-EcoRI/leu2-EcoRI* or *leu2-BstEII*/*leu2-BstEII* gene pair. Therefore, loss of Leu+ frequency can be due to loss of gene conversion, or an increase in outcomes invisible to this assay. The frequency of conversion events is measured by comparing growth on selective and non-selective media. WT strains had a Leu*+* frequency of 1.03+/-0.33×10^-5^ (**Figure 1C and Supplemental Table 3**) that was not significantly different from the *rdh54Δ* strains, which had a conversion frequency of 0.92+/-0.5×10^-5^ (**Figure 1C**). In contrast, the *rdh54T851A* and *rdh54S852A* strains had a significant reduction in Leu*+* frequencies of 0.3+/-0.19×10^-5^ and 0.44+/-0.21×10^-5^, respectively (p<0.0001 and p=0.0004). The two phosphomimic versions of Rdh54, *rdh54T851D* and *rdh54S852D*, were comparable to WT in conversion frequencies (0.9+/-0.15×10^-5^ and 0.88+/- 0.19×10^-5^, respectively) (**Figure 1C**). Surprisingly, the frequency observed for the *rdh54T851A,S852A* double mutant was 1.03+/-0.44×10^-5^ and comparable to WT (**Figure 1C**). This suggests the *rdh54T851A,S852A* allele may be a compensatory mutation. From these data we conclude that *rdh54T851A* and *rdh54S852A*, alter recombination outcomes between homologous chromosomes.

We next asked which sub-population of HR outcomes were altered in mutant *rdh54* backgrounds. When using the homologous chromosome as a template for repair, LOH events can occur by both reciprocal (CO) and non-reciprocal (BIR) exchange of DNA sequences between chromosomes. *RDH54* has previously been shown to reduce BIR outcomes, and to reduce CO frequency when the gene is inactivated by a mutation to the Walker A box, *rdh54K318R* (26,62). We hypothesized that changes to interhomolog gene conversion induced by the *rdh54T851A* and *rdh54S852A* allele may result from an increase in BIR outcomes, or a decrease in CO outcomes resulting in reciprocal exchange. To test this hypothesis, we used a previously described genetic reporter assay that monitors LOH outcomes (45,76). The reporter is based on an *I-Sce1* site located in an inactive copy of the *ade2* gene on budding yeast chromosome XV. The homologous chromosome has a different, but also inactive, allele of *ade2*. Upon *I-Sce1* induction with galactose both sister chromatids are cut, and repair occurs using the homologous chromosome as a template. Each homolog contains a different antibiotic marker located 154 kbp downstream of the break site (**Figure 2A**). Therefore, this assay can differentiate between long tract gene conversion (LTGC), short tract gene conversion (STGC), CO, NCO, and BIR outcomes associated with gene conversion events (45,76) (**Figure 2AB and Supplemental Figure 3AB**).

**Figure 2:**
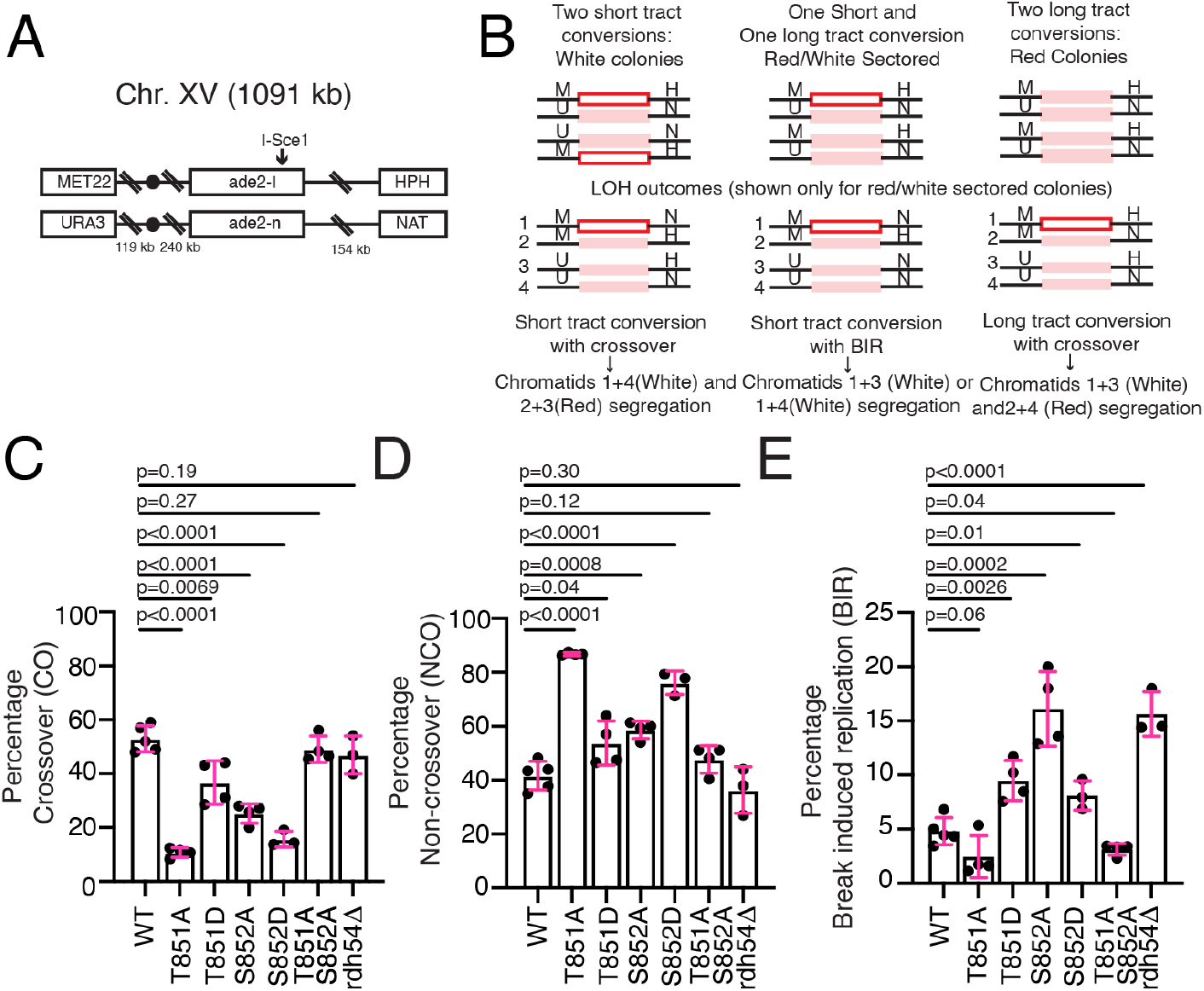
Rdh54 mutants have altered CO to NCO outcomes. (**A).** Schematic diagram illustrating the configuration of the HR reporter assay used in our study to monitor HR outcomes. **(B).** Schematic diagram illustrating the potential outcomes from the induced *I-Sce1* double strand break. The assay can inform on LOH outcomes. **(C).** Percentage of HR outcomes that resulted in CO for WT, *rdh54T851A*, *rdh54T851D*, *rdh54S852A*, *rdh54S852D, rdh54T851A,S852A* and *rdh54Δ*. The bars represent the mean, and the error bars represent the standard deviation for three independent experiments and crosses. (**D).** Percentage of HR outcomes that resulted in NCO for WT, *rdh54T851A*, *rdh54T851D*, *rdh54S852A*, *rdh54S852D, rdh54T851A,S852A,* and *rdh54Δ*. The bars represent the mean, and the error bars represent the standard deviation for three independent experiments and crosses. **(E).** Percentage of HR outcomes that resulted in BIR for WT, *rdh54T851A*, *rdh54T851D*, *rdh54S852A*, *rdh54S852D, rdh54T851A,S852A,* and *rdh54Δ*. The bars represent the mean, and the error bars represent the standard deviation for at least three independent experiments and crosses. Statistical tests were performed using a two tailed t-test.

The clearest determinant of repair outcomes can be seen when one sister is repaired by STGC, and the other by LTGC, resulting in sectored colonies. These colonies have undergone the first chromosomal division after plating and provide the easiest interpretation of the genetic outcomes following the DSB. In our hands sectored colonies result in a near even distribution of CO to NCO outcomes and a small percentage of BIR outcomes. In contrast solid red or solid white colonies result in a majority NCO outcomes (62). This makes it more difficult to determine if there are significant changes in the population of reciprocal and non-reciprocal exchange outcomes. In this study, our specific question was whether there was a change in the ratio of CO:NCO:BIR in Rdh54 mutants. The sectored colonies represent a genetically sensitive population to answer this question. Therefore, we are reporting only outcomes for red/white sectored colonies.

We observed no difference in the ratio of LTGC versus STGC in any of the Rdh54 mutant alleles (**Supplemental Figure 3B**). However, most of the alleles tested did have changes in the distribution of HR outcomes (**Figure 2CDE and Supplemental Table 4**). A general reduction in CO outcomes was observed in *rdh54T851A* (10+/-1.7%, p<0.0001), *rdh54T851D* (36.7+/-8%, p=0.0069), *rdh54S852D* (16.9+/-3.4%, p<0.0001), and *rdh54S852A* (25.2+/-3.6%, p<0.0001) as compared to WT (53+/-10%) (**Figure 2C**). Surprisingly, the *rdh54T851A,S852A* allele was again compensatory and there was no significant difference in CO outcomes relative to the WT. There was also an increase in NCO outcomes associated with most of these alleles. In the case of the *rdh54T851A* allele (86.7+/-0.5%) the magnitude of this change was proportional to the loss of CO outcomes (**Figure 2D**). With *rdh54T851D* (53.7+/-8%, p=0.03), *rdh54S852A* (58.6+/-3.2%, p=0.0008), and *rdh54S852D* (76.1+/-4.2%, p<0.0001) these changes were also accompanied by an increase in BIR outcomes (**Figure 2E**). The magnitude of the increase in BIR was greatest in the *rdh54S852A* strain (16.1+/-3.4%, p<0.0001), with smaller changes in *rdh54T851D* (9.4+/- 1.8%, p=0.0026) and *rdh54S852D* (9.4+/-2.8%, p=0.01). The change in the *rdh54S852A* mutant was comparable to the *rdh54Δ* strains (**Figure 2E**). The *rdh54T851A,S852A* was compensatory in all cases and indistinguishable from WT. From these data we conclude that the T851 and S852 residues in Rdh54 constitutes an important site during HR that can regulate exchange of information between chromosomes.

For BIR events to be observed in the red/white assay a distal antibiotic marker needs to be copied (**Figure 2A**). Therefore, BIR of >150 kbp is required to produce a BIR positive colony. DNA template switching is a DNA repair outcome that is associated with BIR in which the DNA being replicated moves to a site of new homology between chromosomes (70). This is believed to be widespread in meiotic recombination (77) and can occur frequently between homologous chromosomes. Importantly, *RDH54* has been implicated in template switching (70,71). We previously proposed that an increase in BIR observed in a *rdh54Δ* strain could result from a loss in DNA template switching during the BIR reaction. This could be caused by more processive BIR which fails to switch templates more frequently copying the necessary distal antibiotic marker. We reasoned that if this were true then there would also be a loss of template switching associated with the *rdh54S852A* allele.

To test this, we used a previously characterized yeast strain (70) that was designed to test template switching outcomes during BIR by reconstitution of a *URA3* marker located at three distinct sites on chromosome III (70) (**Supplemental Figure 4A**). DSBs were induced by overexpression of the HO nuclease leading to cleavage of a unique HO site. The frequency of Ura+ cells was measured for WT, *rdh54T851A*, *rdh54T851D*, *rdh54S852A*, *rdh54S852D*, and *rdh54T851A,S852A* strains. A Ura+ frequency of 0.45+/-0.13×10^-2^ was measured for the WT strain. This is consistent with previous reports (70) and served as a reference for comparison with the Rdh54 mutants. The *rdh54Δ* strain had a Ura+ frequency of 0.08+/-0.02×10^-2^, a roughly 5.5-fold reduction from WT. There was also significant reduction in Ura+ frequency observed in the *rdh54T851A* strain (0.18+/- 0.02+/-×10^-2^ Ura+, p<0.0001), a 2.5-fold reduction from WT. Surprisingly, there was no significant reduction in template switching observed for *rdh54T851D*, *rdh54S852A*, *rdh54S852D,* or *rdh54T851A,S852A* strains (**Supplemental Figure 4B**). This suggests that the observed increase in BIR associated with *rdh54S852A* is not due to a reduction in template switching activity, and that phosphorylation of the S852 residue in our reporter strain likely limits BIR initiation. We cannot rule out the possibility that the increase in BIR observed in the *rdh54Δ* strains stem from more processive BIR due to loss of template switching, and there is a difference between the null and *rdh54S852A* mutant in this instance, as occurs in other assays. In addition, we also conclude that the *rdh54T851A* allele shows a reduction in template switching that is not as severe as with the *rdh54Δ* strain. This could be caused by a general reduction in BIR outcomes in the *rdh54T851A* strain, as BIR was down relative to WT in the red/white assay but was not deemed statistically significant due to the overall low percentage of BIR outcomes in the WT strain (**Figure 2E**).

### Biochemical analysis of Rdh54 mutants reveal physical changes in activity on dsDNA

To develop a biochemical hypothesis to understand how mutations in Rdh54 may be affecting the activity of Rdh54, we used the Alphafold2 predicted structure of Rdh54 to produce a molecular model of Rdh54 bound to a short fragment of dsDNA (**Figure 3A**). We aligned the Alphafold2 model of Rdh54 to an existing *Sulfolobus solfataricus* Snf2 DNA translocase structure bound to dsDNA (**PDBID:1Z63**) (78). Because the *S. solfataricus* crystal structure represents the open form of the enzyme, we also created a closed model of Rdh54 by rotating one of the RecA domains (**Supplemental Figure 5**) using the procedure from the original study (78). The closed form of the structure represents the active form of the enzyme (**Figure 3A and Supplemental Figure 5**). We were surprised to find that the T851 and S852 residues were located a large distance from the enzyme active site at the end of an alpha-helix predicted to extend along the dsDNA. The modelled structure of the alpha helix has a region of high confidence (90>pLDDT>70, cyan) at the C-terminal end, with less confidence predicted for the interior (70>pLDDT>50, yellow). Rdh54 is known to form an oligomer on dsDNA and the location of these two residues led us to the hypothesis that this region of Rdh54 may affect inter-subunit communication. To test this biochemically, we overexpressed and purified 6xHis-Rdh54 and 6xHis-Rdh54T851A, 6xHis-Rdh54T851D, 6xHis-Rdh54S852A, 6xHis-Rdh54S852D, 6xHis-Rdh54T851AS852A, as well as mCherry versions of each protein from *E. coli* to measure ATP hydrolysis and single molecule activity on dsDNA. The 6xHis tag on this protein does not have detrimental effects on protein activity (28). Throughout this study we will use recombinant 6xHis-Rdh54. For simplicity we will omit using the 6xHis nomenclature in the remainder of the text.

**Figure 3:**
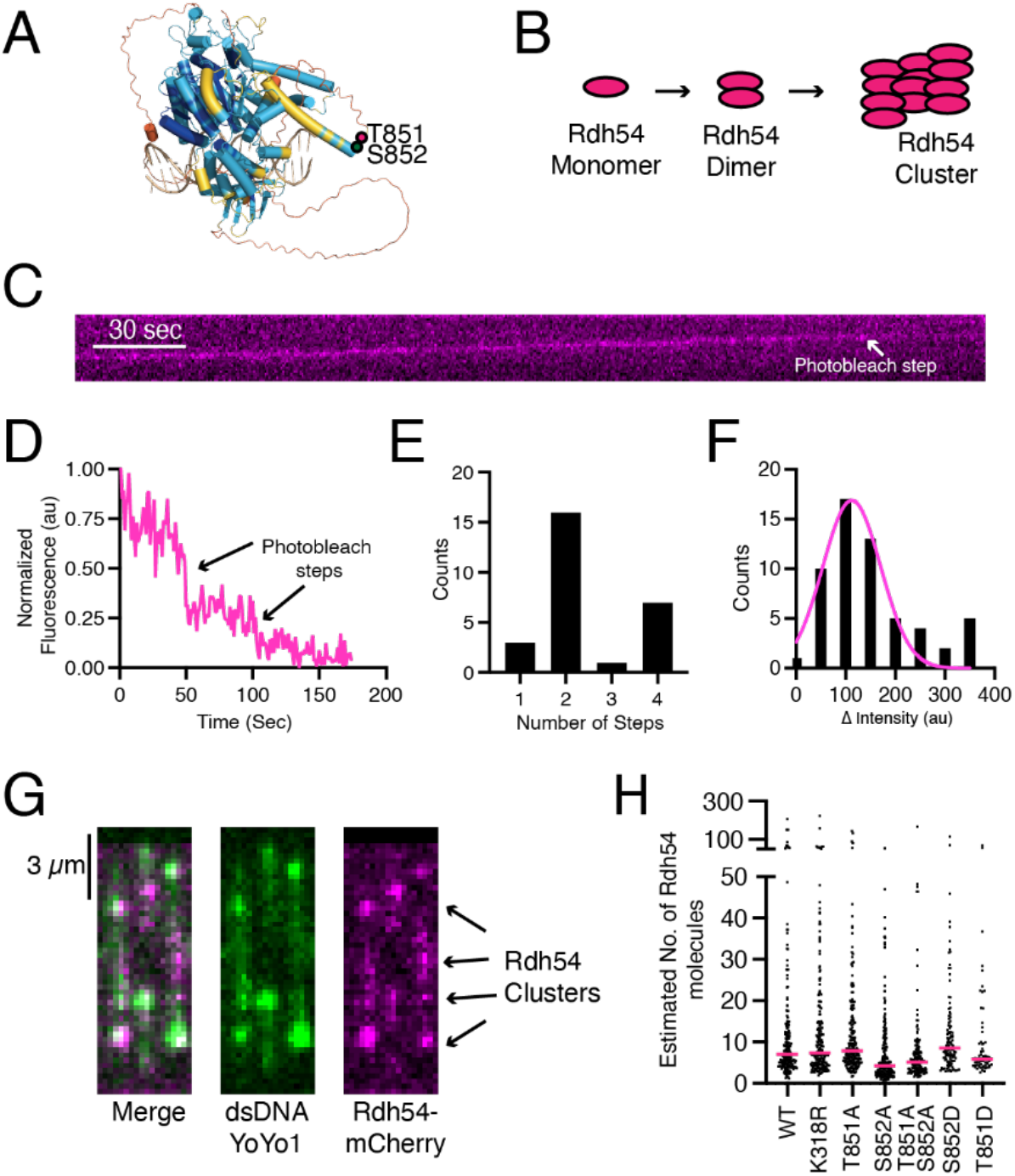
Rdh54 sequentially organizes into clusters on dsDNA. **(A).** AlphaFold2 generated model of Rdh54 bound to dsDNA. The color scheme represents the pLDDT format colored structural model of yRdh54 in a closed conformation. The AlphaFold2 per-residue confidence scores (pLDDT): very high confidence (pLDDT>90), dark blue; high confidence (90>pLDDT>70), cyan; low confidence (70>pLDDT>50), yellow; very low confidence (50>pLDDT), orange. DNA is shown in tan. The location of T851(magenta) and S852 (green) are indicated by dots. The predicted structure was obtained from uniprot, and the model was generated using PDB:1Z63 and PDB:3PJR. **(B).** Cartoon diagram of expected assembly of Rdh54 into clusters. **(C).** Representative kymograph illustrating a translocating Rdh54-mCherry molecule photobleaching. **(D).** Representative trace showing the intensity loss associated with individual photobleaching steps. **(E).** Distribution of photobleaching steps observed for individual Rdh54 molecules bound to the DNA (N=27). **(F).** Distribution of intensity drops associated with photobleaching events. The data are fit by a Gaussian distribution that was used to calculate the mean intensity drop per photobleaching event (N=57). **(G).** Wide field microscopy images of individual dsDNA (YoYo1) molecules bound by Rdh54-mCherry clusters. **(H).** Dot plot representing the distribution of the estimated sizes of Rdh54 clusters for WT (N=201), Rdh54K318R (N=205), Rdh54T851A (N=197), Rdh54S852A (N=214), Rdh54T851AS852A (N=142), Rdh54S852D (N=124), and Rdh54T851D (N=68). The magenta line represents the median of the data. Statistical tests were performed using Mann-Whitney non-parametric tests.

### Rdh54 is a functional oligomer

We used single molecule imaging to directly observe Rdh54-mCherry bound to dsDNA. Biochemically, Rdh54 forms a homo-oligomer both in solution, and when bound to dsDNA (61). To better understand the nature of Rdh54 homo-oligomers bound to dsDNA we used photobleaching measurements to determine the size and distribution of Rdh54 clusters bound to DNA (**Figure 3CD**). For smaller groups of Rdh54-mCherry we counted individual photobleaching steps (**Figure 3E**). The most common size of clusters that could be measured directly contained a dimer of Rdh54 (59%, N=27) (**Figure 3E**). The second most common (26%, N=27) group contained four Rdh54 molecules. Groups of one and three units were also observed but occurred less frequently than the other groups (**Figure 3E**). The majority of Rdh54 formed larger clusters on the DNA that could not be directly measured by photobleaching. To estimate the size of these clusters, we measured the intensity drop associated with individual photobleaching event and created a distribution from this data. The distribution was used to generate a mean intensity for an individual fluorophore (**Figure 3F**). We then divided the measured intensities for Rdh54 clusters bound to DNA by the mean intensity to create an estimated number of Rdh54-mCherry molecules per cluster (**Figure 3GH**). These measurements resulted in a large distribution of cluster sizes with a median value of 7.06 molecules per cluster for WT-Rdh54 (N=201). We also measured cluster sizes for each of the Rdh54 mutants. We found there was no significant difference in cluster sizes between WT, and Rdh54T851A (p=0.14), Rdh54T851D (p=0.28), Rdh54S852D (p=0.08), and Rdh54K318R (p=0.38) (**Figure 3H**). In contrast, Rdh54S852A and Rdh54T851AS852A were significantly different, with median cluster sizes of 4.22 (p<0.0001) and 5.1 (p<0.0001), respectively (**Figure 3H**). From this data we conclude that Rdh54 forms clusters on dsDNA and that Rdh54S852A and Rdh54T851AS852A have reduced clustering activity.

To determine if clustering had functional consequences, we first determined if different populations of Rdh54 could mix within clusters by combining Rdh54-mCherry and Rdh54-GFP and directly visualizing the outcome via single molecule imaging. In these experiments we measured the percentage of co-localization and the translocation velocity along DNA. Rdh54-mCherry colocalized with Rdh54-GFP in 93% of cases (**Supplemental Figure 6AB**, n=132/142) indicating an interaction between the two populations of molecules. The mixed molecules co-translocated with a mean velocity of 68.5+/-40.4 bp/sec, a mean distance of 14.3+/-8.8 kbp **(Supplemental Figure 6CD**) and were indistinguishable from un-mixed Rdh54-mCherry (N=32 and N=32, p=0.13 and p=0.22, respectively). We next performed classic translocase poisoning experiments where we mixed an equal amount of WT protein with either Rdh54K318R or Rdh54T851A. The hypothesis was that if proteins within a cluster were functionally dependent on each other, then incorporating mutant versions of the protein would poison the WT protein and reduce the velocity of the cluster. When we mixed an equal molar amount of Rdh54-mCherry with unlabeled Rdh54K318R, the mean velocity of the total population was reduced to 30.7+/-20 bp/sec (N=62, p=<0.0001), representing a significant change from WT (**Supplemental Figure 6CD**). This is consistent with interdependency of Rdh54 subunits with the cluster. We also observed reduced rates of translocation (23+/-16 bp/sec, N=31, p=<0.0001) (**Supplementa**l **Figure 6CD**) when unlabeled Rdh54T851A was mixed with equimolar amounts of WT. These data support an oligomeric motor configuration of Rdh54 in which translocation is contingent on the composition of the cluster.

Next, we reasoned that if cluster formation had an impact on HR outcomes, then strains that were heterozygous for *rdh54K318R* and *rdh54T851A* alleles would have altered CO/NCO ratios. We chose to test CO/NCO ratios because this was the most significant difference we observed in our initial experiments and would likely be sensitive enough to detect even a partial change in outcomes. We performed red/white recombination experiments with *RDH54:rdh54K318R* and *RDH54:rdh54T851A* strains. We found that the *RDH54:rdh54K318R* had a reduction in CO (27.3+/-4.0% versus 53 +/-10%, p=0.005) in favor of NCO (67.9+/-6% versus 41+/-10%, p=0.004) outcomes. Likewise, there was a reduction in CO (34+/-9% versus 53+/-10%, p=0.002) and an increase in NCO outcomes (64.2+/-7% versus 41+/-10%, p=0.003) in the *RDH54:rdh54T851A* strain (**Supplemental Figure 6E**). From these data we conclude that Rdh54 forms biologically active clusters during DNA damage response that are likely modulated by inter-subunit communication.

### Mutations in Rdh54 result in de-coupling of ATP hydrolysis from translocation

Next, we measured the biochemical activity of the Rdh54 mutants. We initially measured the ATP hydrolysis activity at 100 µM and 1000µM ATP concentrations (**Figure 4A**). In both cases we found that WT and Rdh54T851D produced the same amount of ADP at all time points measured. In contrast, the Rdh54S852A mutant produced less ADP indicating a defect in ATP hydrolysis. The opposite effect was observed for the Rdh54T851A which produced higher levels of ADP than WT at all time points (**Figure 4A**). Like the Rdh54T851A mutant, the Rdh54S852D and Rdh54T851AS852A mutants produced a greater amount of ADP over the time course. However, this difference was not as severe as that observed for the Rdh54T851A mutant (**Figure 4A**). From this data we conclude that mutations in this region of Rdh54 alter ATP hydrolysis activity.

**Figure 4:**
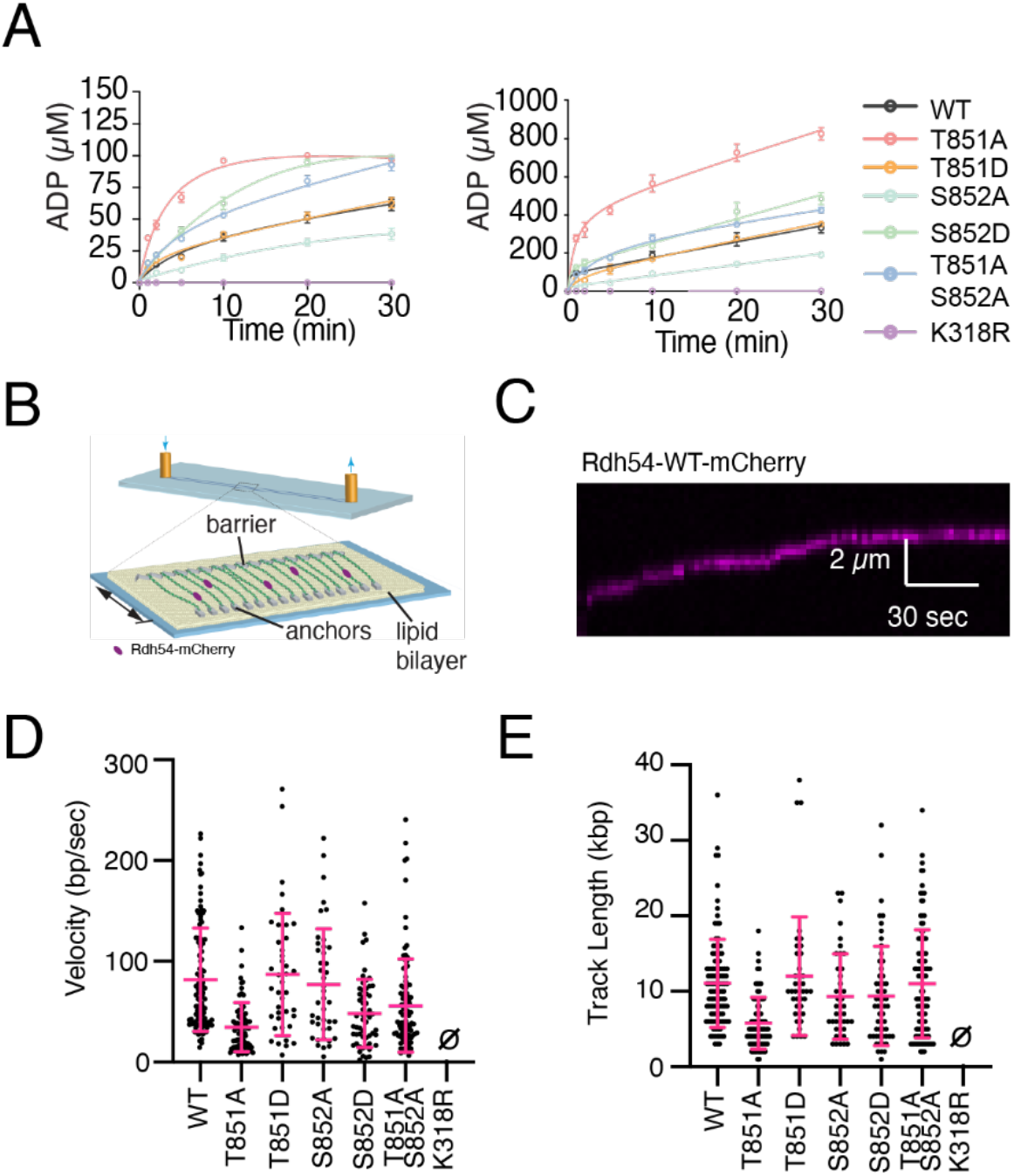
Mutations in Rdh54 result decoupling of ATP hydrolysis from translocation. **(A).** ATP hydrolysis activity for Rdh54, Rdh54T851A, Rdh54T851D, Rdh54S852A, Rdh54S852D, Rdh54T851AS852A, and Rdh54K318R at a total ATP concentration of 100 µM (Left) and 1000 µM (Right). The points equal the mean, and the error bars the standard deviation of three independent experiments. **(B).** Schematic diagram illustrating DNA curtains setup, a single molecule approach that allows measurement of Rdh54-mCherry velocity and track length. **(C).** Representative kymograph illustrating the translocation of Rdh54-mCherry along dsDNA. **(D).** Translocation velocities (bp/sec) for individual translocation events of Rdh54 (N=110), Rdh54T851A (N=81), Rdh54T851D (N=40), Rdh54S852A (N=42), Rdh54S852D (N=56), Rdh54T851AS852A (N=101), and Rdh54K318R. The crossbar represents the mean of the data, and the error bar represents the standard deviation of the experiment. **(E).** Track lengths (kbp) for individual translocation events of Rdh54 (N=110), Rdh54T851A (N=81), Rdh54T851D (N=40), Rdh54S852A (N=42), Rdh54S852D (N=56), Rdh54T851AS852A (N=101), and Rdh54K318R. The crossbar represents the mean of the data, and the error bars represent the standard deviation of the data.

We used single molecule imaging (**Figure 4B**), to measure the movement of Rdh54 along dsDNA (**Figure 4BC**). For WT Rdh54-mCherry, we measured a translocation velocity of 81.7+/-51 bp/sec (N=110), and a track length of 12.1+/-7.1 kbp (N=110) (**Figure 4DE**). Both observations are consistent with previous measurements made in different laboratories (60,61). For the Rdh54T851A mutant we observed a reduction in both mean velocity (35.4+/-26 bp/sec, N=81, p=<0.0001) and mean track length (5.1+/-3.0 kbp, N=81, p<0.0001). In contrast, the Rdh54T851D mutant was not different from WT in either mean velocity (86+/-60 bp/sec, N=40, p=0.877) or mean track length (12.0+/-7.8 kbp, N=40, p=0.9), consistent with the results from the ATP hydrolysis experiments. The reduction in velocity of the Rdh54T851A mutant and the increase in ATP hydrolysis is consistent with a less efficient coupling of ATP hydrolysis to translocation. The Rdh54S852A mutant translocated with a mean velocity of 77+/-54 bp/sec (N=41) and a mean track length of 9.3+/-5.6 kbp (N=41). These values were not significantly different from the WT and suggest a mutant which is more efficient at coupling ATP hydrolysis to translocation. In contrast, Rdh54S852D had a significant decrease in velocity (48.3+/-33.3 bp/sec, N=60, p<0.0001) and a small decrease in track length (9.3+/-6.5 kbp, N=60, p=0.03) relative to WT (**Figure 4DE**). When combined with the ATP hydrolysis results, the S852A and S852D mutants have inverse behaviors relative to WT, and this change is consistent with altered coupling of ATP hydrolysis to translocation. Finally, we tested the activity of the Rdh54T851AS852A mutant. We found this mutant had higher mean velocity (55+/-46 bp/sec) than the Rdh54T851A single mutant but was still significantly less than WT (p<0.0001, N=101). We saw an insignificant difference in track length (11.0+/-7.1 kbp, p=0.34) (**Figure 4DE**) compared to the WT. This mutant was interesting because it has translocation defects like the Rdh54T851A mutant, but it has reduced clustering capacity like the Rdh54S852A mutant. These data suggest that the compensatory phenotypes observed in cells are an averaging effect caused by defects associated with each individual mutant. Together our data suggests that mutations in this region of Rdh54 lead to altered biochemical activity on dsDNA that may be due to changes in inter-subunit communication.

### Rad53 forms a physical interaction with Rdh54

The Rdh54S852 residue is part of a consensus Rad53 phosphorylation site, and previous reports genetically link the kinase Rad53 to phosphorylation of Rdh54 (59). However, a direct interaction between these proteins has not been observed. Rad53 is an auto-kinase that can phosphorylate copies of itself in trans (49). Because of this, it can be purified in a hyper-phosphorylated form from *E. coli* (52,79,80). Likewise, a single point mutant in Rad53, Rad53D339A, can eliminate kinase function and results in the purification of an inactive form of Rad53. To directly observe the interaction between Rdh54 and Rad53 we generated GFP-Rad53 and GFP-Rad53D339A constructs and purified them from *E. coli*. The GFP-Rad53 retains auto kinase function and can be purified in a phosphorylated form (**Supplemental Figure 7A**). We measured the interactions between GFP-Rad53/Rdh54-mCherry and observed direct and stable binding via single molecule imaging (**Figure 5A**). We next asked if there was an effect on Rdh54 cluster size in the presence of Rad53. We were surprised to find that the median estimated size of Rdh54 clusters was slightly larger at 0.1 nM Rad53 (7.0 versus 8.6, p=0.003), and twice as large at 10 nM Rad53 (7.0 versus 15.5, p<0.0001), suggesting that Rad53 may affect Rdh54 cluster size (**Figure 5B**). To determine if this was dependent on phosphorylation of Rad53 we measured the interaction between Rdh54-mCherry and GFP-Rad53D339A. We also observed a stable binding interaction between this pair (**Figure 5A**). We found that the effect of 10 nM GFP-Rad53D339A on Rdh54 cluster size was reduced (15.5 versus 10, p<0.0001) (**Figure 5C**). This also corresponded with a reduced intensity of GFP-Rad53D339A bound to Rdh54 (**Figure 5D**) and a reduced percentage of Rdh54-mCherry clusters that were occupied by GFP-Rad53D339A relative to WT (**Figure 5E**). Finally, we observed a strong correlation between the signal intensity of Rdh54-mCherry and GFP-Rad53 in the WT (r=0.7), but only a weak correlation with the GFP-Rad53D339A mutant (r=0.2, **Figure 5F**). This suggests that more molecules of Rdh54 per cluster resulted in more molecules of Rad53 binding. It should be noted that the inclusion of 10 nM GFP-Rad53D339A still resulted in larger clusters than Rdh54 alone. From these data we conclude that Rdh54 and Rad53 form an interaction that is independent of Rad53 phosphorylation. However, phosphorylation of Rad53 enhances the interaction with Rdh54. These data also identify a phosphorylation independent role of Rad53 in the growth of Rdh54 clusters.

**Figure 5:**
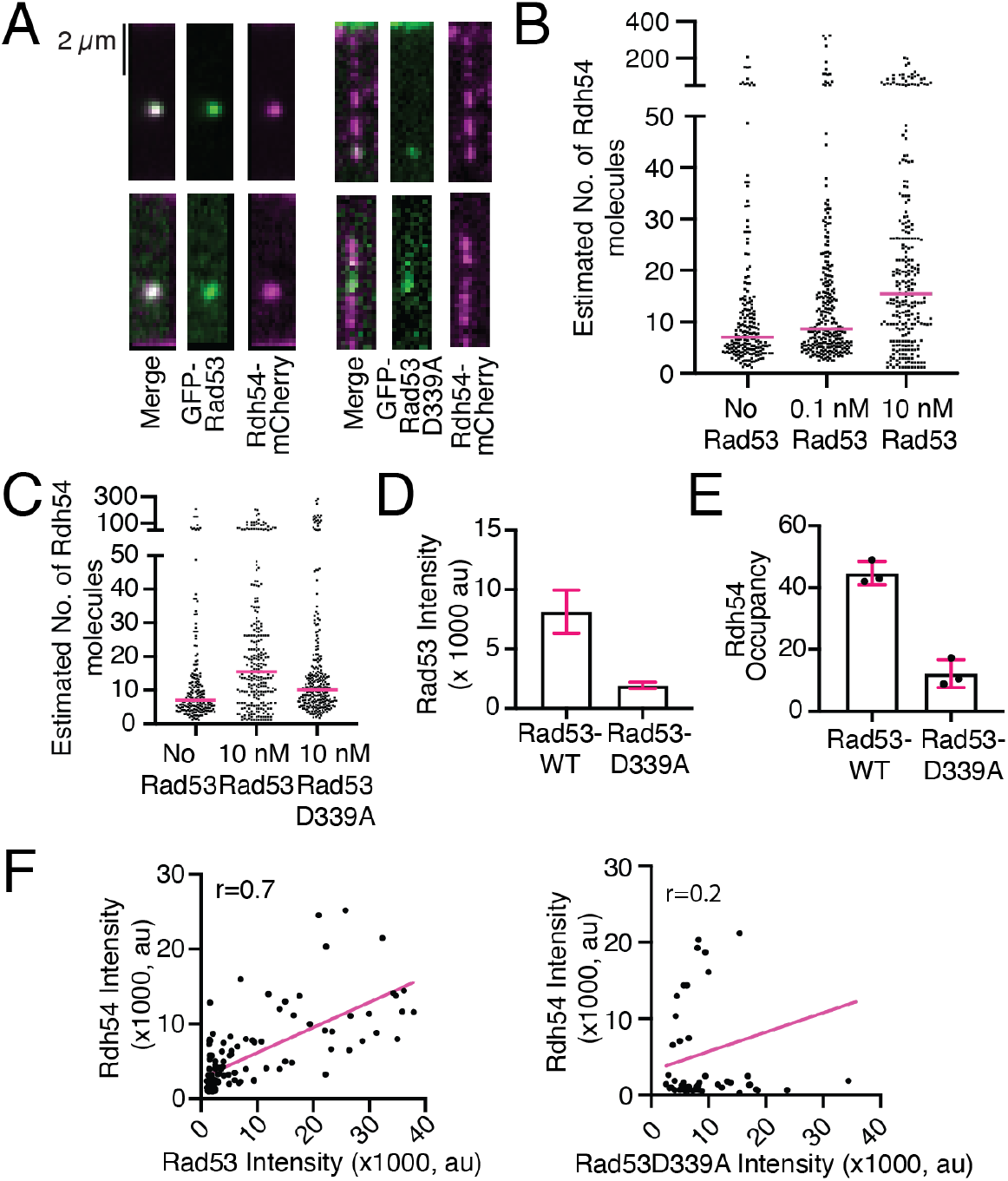
Phosphorylated Rad53 preferentially interacts with Rdh54 clusters. **(A).** Widefield TIRF microscope images of individual DNA molecules bound by Rdh54-mCherry with GFP-Rad53 (left) and Rdh54-mCherry with GFP-Rad53D339A (Right). **(B).** Dot plot representing the distribution of estimated sizes of Rdh54-mCherry clusters bound to DNA with no GFP-Rad53 (N=201), 0.1 nM GFP-Rad53 (N=273), and 10 nM GFP-Rad53 (N=262). The crossbar represents the median of the data. **(C).** Dot plot representing the distribution of estimated sizes of Rdh54-mCherry clusters bound to DNA in a population. Including No GFP-Rad53 (N=201), 10 nM GFP-Rad53 (N=262) and 10 nM GFP-Rad53D339A (N=282). The magenta line equals the median of the data. **(D).** Graph representing the signal intensity for GFP-Rad53 and GFP-Rad53D339A bound to Rdh54-mCherry. The bar represents the mean of the data, and the error bars represent the 95% confidence interval of the distribution. **(E).** A graph representing the percentages of Rdh54-mCherry clusters occupied by GFP-Rad53 and GFP-Rad53D339A. The bar represents the mean, and the error bar represents the standard deviation of at least three independent experiments. **(F).** Graph representing the correlation of intensity measurements for Rdh54-mCherry with Rad53-WT. The WT in B and C is reproduced from Figure 3.

We next evaluated whether there was an *in situ* interaction between Rad53 and Rdh54 using a co-immunoprecipitation experiment. We used a strain of yeast with an HO inducible break that lacked a donor DNA sequence. We used this strain because it had previously been used to demonstrate a genetic interaction between Rad53 and Rdh54 (59). We found that there was an *in-situ* interaction between Rad53 and Rdh54, as predicted by our biochemical experiments (**Supplemental Figure 8 and Figure 6A**). This interaction was not dependent on DSB formation and formed both with and without galactose. We also tested the interaction between *rdh54T851A* or *rdh54S852A* and Rad53. We found that these mutants were able to interact with Rad53 in a galactose independent manner (**Figure 6A**). From these data we conclude that Rad53 and Rdh54 interact *in situ*. Our co-immunoprecipitation experiments are non-quantitative, so we next evaluated the interaction between Rad53 and Rdh54T851A or Rdh54S852A *in vitro* using single molecule imaging. Not surprisingly, both proteins interacted with Rad53 (**Figure 6C**). For the Rdh54T851A mutant the resulting change in Rdh54 cluster size was comparable to WT (15.5 versus 12.1, p=0.50). In contrast, the Rdh54S852A mutant had reduced cluster size (5.7 versus 15.5, p<0.0001) (**Figure 6C**). This was comparable to the difference between WT-Rdh54 and the Rdh54S852A mutant in the absence of Rad53. However, we did observe that fewer clusters of the Rdh54S852A mutant were bound by Rad53 (**Figure 6D**). This was contrasted by the Rdh54T851A mutant which had more clusters with Rad53 bound than WT (**Figure 6D**). From these data we conclude that mutations in Rdh54 lead to an altered interaction with Rad53.

**Figure 6:**
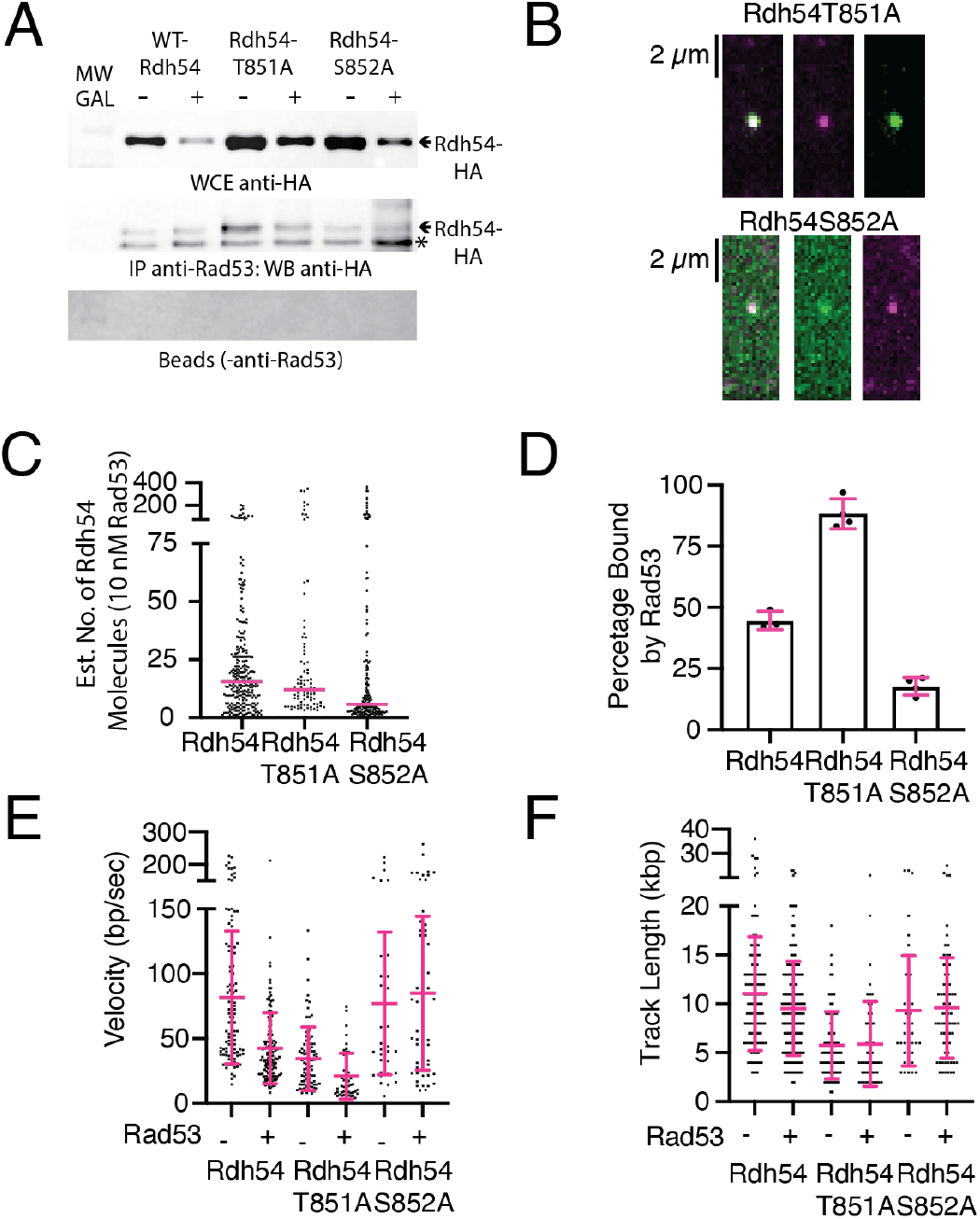
Interaction between Rad53 and Rdh54 affects translocation rates. **(A).** Co-IP experiments showing an *in situ* interaction between Rdh54 mutants and Rad53. Shown are WCE (Top), IP samples (Middle), and beads without antibodies (Bottom). * Is an unspecific band that also appears in the No HA tagged (Supplemental Figure 8) **(B).**Widefield TIRF microscope images illustrating the interaction between Rdh54T851A-mCherry (magenta) and GFP-Rad53 (green), and the interaction between Rdh54S852A-mCherry (magenta) and GFP-Rad53 (green). **(C).** Dot plot representing the estimated number of Rdh54 molecules in each cluster for WT (N=262), Rdh54T851A-mCherry (N=88), and Rdh54S852A-mCherry (N=167) in the presence of 10 nM GFP-Rad53. The crossbar represents the median of the data. **(D).** Graph representing the percentage of Rdh54 clusters occupied by GFP-Rad53 for Rdh54-mCherry (N=3), Rdh54T851A-mCherry (N=4), and Rdh54S852A-mCherry (N=4). The bars represent the mean, and the error bars represent the standard deviation of at least 3 independent experiments. The data for Rdh54-mCherry is reproduced from Figure 5. **(E).** Dot plot representing the velocity measurements for Rdh54-mCherry without (N=110) and with Rad53 (N=142), Rdh54T851A-mCherry without (N=81) and with Rad53 (N=49), and Rdh54S852A-mCherry without (N=42) and with (N=66) Rad53. The crossbar represents the mean of the data, and the error bars represent the standard deviation. **(F).** Dot plot representing the track length measurements for Rdh54-mCherry without (N=110) and with Rad53 (N=142), Rdh54T851A-mCherry without (N=81) and with Rad53 (N=49), and Rdh54S852A-mCherry without (N=42) and with (N=66) Rad53. The crossbar represents the mean of the data, and the error bars represent the standard deviation. In E and F, the (-) condition is reproduced from Figure 3.

We next measured the ability of Rad53 to phosphorylate Rdh54 using an *in vitro* kinase assay. We found that Rad53 was able to directly phosphorylate Rdh54 *in vitro* (**Supplemental Figure 7B**). To further validate that phosphorylation was occurring at the S852 site we performed *in vitro* phosphorylation with mass spectrometry. When we measured the recombinant Rdh54 alone we only observed a single phosphorylation mark both in the absence and presence of ATP (**Supplemental Table 5**), and this protein was not phosphorylated at S852. Likewise, only a single phosphorylation mark was observed when Rdh54 was treated with Rad53D339A (**Supplemental Table 5**). In contrast, WT-Rad53 phosphorylated Rdh54 at many sites. Many of these sites are likely non-specific because they were not identified by the Superphos database (**Supplemental Figure 7C and Supplemental Table 5**). However, we did identify four residues that were phosphorylated in both the SuperPhos database and in our *in vitro* assay. These sites included T851 and S852 (**Supplemental Table 7C and Supplemental Table 5**). It should be noted that it is difficult to separate two adjacent potential phosphorylation sites via mass spectrometry, and the phosphorylation mark could be on T851 or S852. Based on these data we conclude that this region is being phosphorylated both *in vivo* and *in vitro*.

Our single molecule data suggested that the phosphomimic mutant of S852, S852D was slow to translocate on DNA. We reasoned that we could directly test whether phosphorylation of the S852 residue caused a similar defect in translocation by incubating Rdh54, Rdh54T851A, or Rdh54S852A with catalytic amounts of Rad53 and monitoring translocation activity. By using phosphorylation site mutants of Rdh54 we could reduce the contribution of non-specific phosphorylation and focus on sites that were identified both *in vitro*, *in vivo*, and had phenotypic consequences. We therefore incubated Rdh54-mCherry, Rdh54T851A-mCherry, or Rdh54S852A-mCherry with catalytic amounts of Rad53 for 60 minutes prior to measuring the translocation velocity via single molecule imaging. Rdh54-mCherry moved with a velocity of 42+/-21 bp/sec (**Figure 6E**, N=140). This was significantly (p<0.0001) different from the translocation velocity of 81.7+/-51 bp/sec observed for untreated Rdh54-mCherry, and comparable to the translocation velocity observed for Rdh54S852D (48+/-33.7 bp/sec). The translocation length for Rad53-treated Rdh54-mCherry (9.5+/-4.5 kbp) (**Figure 6F**, N=140) was only slightly lower than that observed for WT and was the same as Rdh54S852D. When we measured the velocities of Rdh54T851A-mCherry we found a reduction in velocity to 20+/-17 bp/sec as compared to Rdh54T851A-mCherry untreated (34+/-24 bp/sec, p<0.0001) (**Figure 6E**). There was no difference in the track length (**Figure 6F**). This is consistent with the T851 site not being the kinase site. However, when we measured the treated Rdh54S852A-mCherry we observed no difference in the translocation velocity along dsDNA (84.9+/-57 bp/sec, N=66 versus 77+/-54 bp/sec, N=42, p=0.48) (**Figure 6E**), or track length on dsDNA as compared to untreated Rdh54S852A-mCherry (**Figure 6F**). Together these data suggest that phosphorylation of Rdh54 by Rad53 at the S852 site slows Rdh54 translocation velocity on dsDNA.

## Discussion

In this study we investigated the regulation of the Rdh54 motor protein during homologous recombination. We identified a previously uncharacterized region of Rdh54 that is critical for function during HR. Mutations to this region caused changes in Rdh54 cluster formation on dsDNA, translocation along dsDNA, and interactions with the effector kinase Rad53. This region is a target of Rad53 whose phosphorylation of Rdh54 results in a reduction in translocation velocity on dsDNA. Importantly, mutants in this region have phenotypic consequences that result in changes in the distribution of LOH outcomes in yeast reporter strains. Based on these findings we present a model where the association and dissociation of Rdh54 clusters from HR repair foci are important for directing HR outcomes.

### Mutations in the C-terminal region of Rdh54 alter structure and function

It has been known for several years that Rdh54 is recruited to HR foci by Rad51. This occurs through an interaction between Rad51 and the disordered N-terminal region of Rdh54 (28,30,81). The interaction is highly cooperative, and self-association of Rdh54 molecules likely contributes to its own recruitment (28). In our experiments we defined the size and distribution of self-associating Rdh54 clusters on dsDNA and linked this activity to genetic consequences. While most of the clustering activity is still likely to occur through the N-terminal domain, a single point mutant in the C-terminal region of the protein, S852A, led to a reduction in cluster sizes bound to dsDNA. Enzymatically Rdh54S852A was more efficient at coupling ATP hydrolysis to translocation on dsDNA. In contrast, conversion of this residue to aspartic acid (S852D), led to no changes in cluster size, but instead de-coupled ATP hydrolysis from translocation. We interpret these results to mean that the S852 residue in Rdh54 is regulating the coupling of ATP hydrolysis to translocation. Its connection to cluster size and ATP hydrolysis likely indicates that this interaction occurs in trans between adjacent subunits of Rdh54. By extension, this suggests that phosphorylation of this residue can regulate the crosstalk between adjacent Rdh54 subunits leading to an overall reduction in translocation velocity on DNA. It is unclear if phosphorylation also results from decoupling of ATP hydrolysis or if it alters the interaction between subunits.

When mutated to alanine the T851 residue has the most severe defect both *in vivo* and *in vitro* caused by significant de-coupling of ATP hydrolysis from translocation on dsDNA. There was no difference in Rdh54 cluster size in this mutant. In contrast, the phosphomimic version of this residue, T851D, had no discernable biochemical defect. Without a high-resolution structure it is difficult to determine how these mutations regulate cluster size and ATP hydrolysis rates. However, an attractive model is that Rdh54 may behave like other RecA proteins in forming a short filament along dsDNA. In this case adjacent protomers may form a shared active site like that observed in Rad51. There is evidence that Rad51 can be phosphorylated at the protomer-protomer interface which prevents binding to dsDNA (82). A similar mechanism may reduce translocation by regulating the coupling of ATP hydrolysis to movement. More work will be needed to test this model.

Phenotypically mutations at T851 and S852 result in different outcomes during interhomolog recombination. We initially tested these mutants in an assay that measures interhomolog recombination between heteroalleles without induction of a break. Alanine substitutions at 851 and 852 sites caused reduced leu+ phenotypes. To determine the cause of these loses, we used a reporter assay that could differentiate between reciprocal and non-reciprocal exchange of information between chromosomes. We found that the T851A mutant had lost reciprocal exchange outcomes due to increases in NCO outcomes at the expense of CO outcomes. In contrast, BIR increased in the S852A mutant at the expense CO outcomes. These outcomes are consistent with a loss of leu+ gene conversion by a reduction in reciprocal exchange or an increase in BIR. We observed that S852D resulted in a reduction in CO outcomes but did not result in a reduction in leu+ conversion frequency. We believe that this is due to differences in the assays with the red/white assay potentially being more sensitive to smaller changes in protein function. For example, S852D still translocates along dsDNA at a higher rate than the T851A mutant, and therefore it is not surprising phenotypes associated with this mutant may be less severe. Likewise, we were unable to see any biochemical differences between Rdh54 and Rdh54T851D, and still saw a slight increase in BIR outcomes and increased MMS sensitivity. One possible reason for this is an alteration in the phosphorylation activity of the adjacent S852 residue. Alternatively, changes in this residue may alter Rdh54 oligomer structure or alter interactions with other proteins during HR based repair. It is also possible that the MMS phenotype is not truly reflective of HR repair outcomes and may represent a different function of Rdh54. If any of these outcomes are true, it does not alter our overall interpretation of the data.

Based on these data we propose a model in which the kinetic lifetime of Rdh54 at DNA repair foci helps determine recombination outcomes. The lifetime is controlled by the on-rate (k_on_), which is a combination of recruitment of Rdh54 by Rad51and the self-association of Rdh54 molecules, and the off-rate (k_off_), which is determined by how quickly Rdh54 translocates away from HR repair intermediates on dsDNA (**Figure 7A**). The off rate is regulated by Rad53 (see below). Based on our data this allows us to group our mutants into categories with the T851A and S852D mutants representing fast clustering mutants. The off rate for these mutants is significantly slower than WT and this results in fast clustering of Rdh54 and an increase in NCO outcomes during interhomolog recombination (**Figure 7B**). Likewise, the S852A mutant represents a mutant that is slow to cluster due to diminished cluster size and increased coupling of ATP hydrolysis to translocation. As discussed below, this mutant cannot be phosphorylated and that will prevent down regulation of translocation along dsDNA. This mutant will have a slow on-rate and fast off-rate leading to an increase in BIR, as is the case in the *rdh54Δ* where no clusters form (**Figure 7B**).

**Figure 7:**
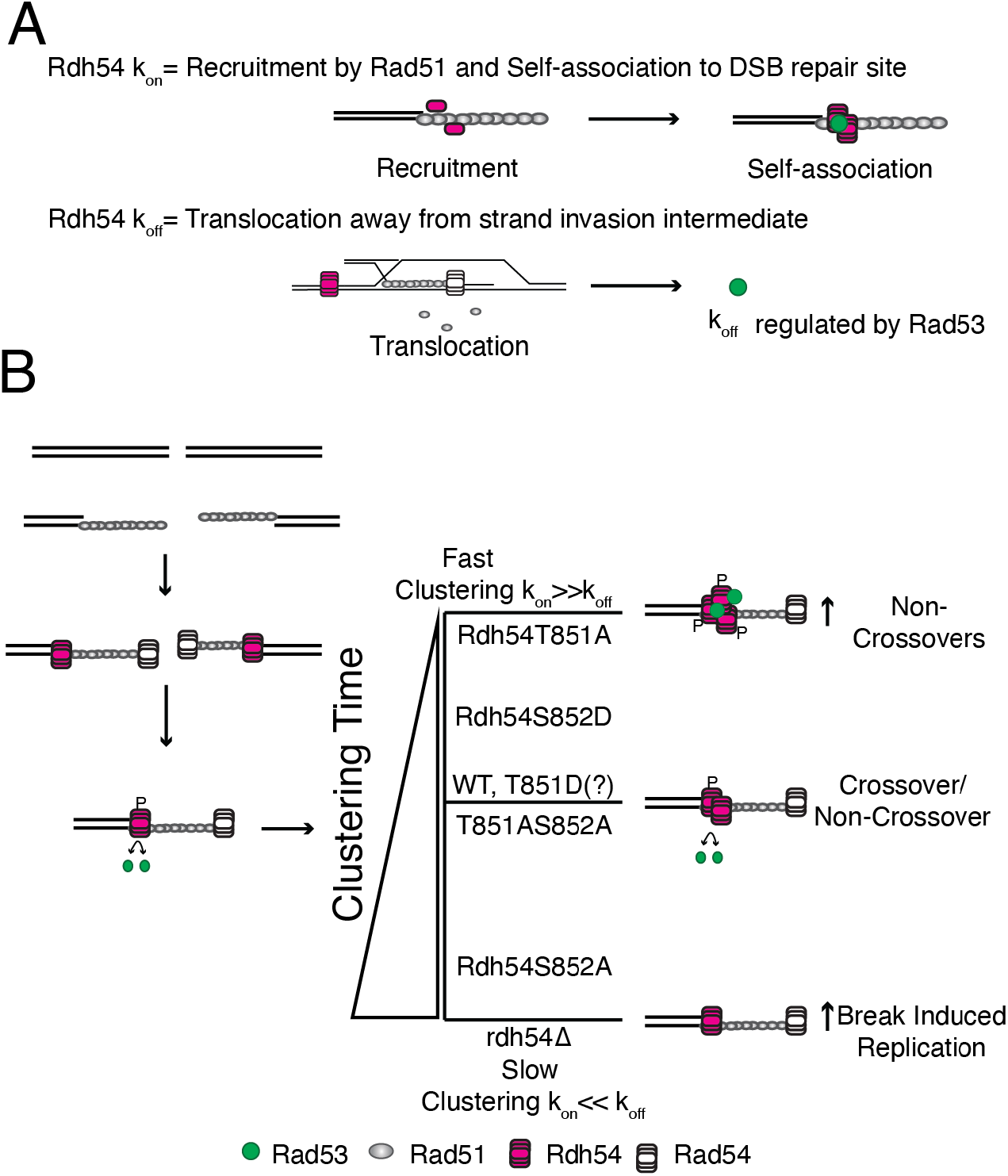
Model for the role of Rdh54 clustering in homologous recombination. **(A).** A model illustrating components in the lifetime of Rdh54 at HR repair foci. The recruitment is initiated by Rad51, and there are additional contributions from the self-association of Rdh54. This is defined as the k_on_. The k_off_ is defined as the rate of translocation away from strand invasion intermediates that occur during recombination. This step is regulated by Rad53 phosphorylation. Rad53 can also alter Rdh54 cluster size. **(B).** Model for biological output of Rdh54 clustering during HR. Our model suggests that fast clustering leads to the development of non-crossover outcomes. In contrast, slow clustering leads to increases in break induced replication. The rate of cluster formation is controlled by the on/off rate of the Rdh54 translocase from HR intermediates.

Importantly, we found that the *rdh54T851A,S852A* allele restored WT function in our genetic assays. Biochemically, this mutant exhibited reduced cluster formation, and a reduction in translocation velocity along DNA. However, this mutant was slightly better at cluster formation than the S852A mutant, and it was better at coupling hydrolysis to translocation than the T851A mutant. Finally, it cannot be phosphorylated due to the mutation at the S852 site. This leads to a combination of variables that likely average out to yield the equivalent of a WT protein *in vivo*. Put another way, the mutant protein likely has the same lifetime as the WT protein at HR repair foci (**Figure 7B**).

### Rdh54 is directly connected to Rad53

Rdh54 is a target of the Mec1/Rad53 signaling axis (59), and physically interacts with the effector kinase Rad53. Once activated during the DNA damage response Rad53 phosphorylates itself through an interaction with Mrc1 at replication forks or through interactions with the Rad9 scaffold (46,53). Activated Rad53 can diffuse around the nuclear space and phosphorylates targets that slow down active repair processes to ensure genomic integrity. We were surprised to observe that Rdh54 formed a stable, direct interaction with Rad53. This interaction was not dependent on phosphorylation, but phosphorylated Rad53 did bind with higher frequency. Likewise, there was an increase in Rdh54 cluster size in the presence of Rad53. We observed that Rad53 preferentially interacted with larger clusters of Rdh54 and had a reduced interaction with Rdh54S852A. Together these data suggest that Rad53 may interact with Rdh54 through a complex set of interactions instead of at a specific domain interface. Functionally, site specific phosphorylation of Rdh54S852 slows the translocation velocity of Rdh54 along dsDNA, and a protein-protein interaction may physically enhance cluster size leading to two separate ways Rad53 can regulate Rdh54 cluster size or growth: a physical interaction that is phosphorylation independent, and downregulation of translocation activity which alters Rdh54’s off-rate that is phosphorylation dependent.

It has previously been shown that Rad53 has the opposite effect on the size of Rad9 clusters in which phosphorylation of the Rad9-BRCT domain leads to reduction in Rad9 cluster size (83). It has been proposed that Rad53 activity may aid in coordinated reduction in Rad9 cluster size. Interestingly, *rad9Δ* strains exhibit a reduction in BIR (84). The opposite is true for Rdh54, and our model would predict that Rad53 may help grow Rdh54 cluster size through both enzymatic and non-enzymatic activities. This counteracts the occurrence of BIR. Larger clusters of Rdh54 may also help to trap Rad53 within a local environment reducing free diffusion, enhancing both Rad53 auto-phosphorylation as well as phosphorylation of Rdh54 and/or other proteins that may occupy similar three-dimensional space. This would create a feed forward mechanism where phosphorylation increases Rdh54 cluster size which could interact with more molecules of Rad53 that then undergo autophosphorylation. It has previously been shown that *rdh54K318R* leads to an accumulation of hyperphosphorylated Rad53 in yeast strains that lack a homologous donor sequence (59), suggesting that Rdh54 clusters may contribute to signal maintenance in this context. It is unclear what would counteract cluster growth and there is likely a missing component to this reaction that leads to diminishing cluster size once DNA repair has occurred. Interestingly, Rad54 has recently been implicated in the development and growth of Rad54 foci in cells, and it is possible that Rdh54 and Rad54 cluster growth is related as they can co-occupy Rad51 filaments during HR mediated repair (85). Ultimately, the interaction between Rad53 and Rdh54 appears to be important for regulation of HR outcomes.

## Conclusion

Here we present a model where the lifetime of Rdh54 at HR foci helps regulate genetic outcomes. In the *S. cerevisiae* model cluster formation is regulated by specific protein-protein interactions within Rdh54 and through an interaction with Rad53. However, this specific interaction may not be conserved in higher eukaryotes. The homolog of Rdh54 in higher eukaryotes is RAD54B. This homologous motor protein may regulate recombination outcomes through a similar clustering type mechanism that controls the on and off rate of this motor protein at HR repair centers. Thus, even if specific interactions are not conserved, regulation is likely to be conceptually the same or at very least similar. Here we present a new model for the recruitment and regulation of Rdh54 during mitotic recombination between homologous chromosomes.

## Data Availability Statement

The data in this manuscript is available upon request.

## Acknowledgements

We would like to acknowledge Hannah Klein, Jim Haber, and Lorraine Symington for yeast strains used in this study. We would also like to thank Dirk Remus for the kind gift of the Rad53 overexpression plasmid. We would also like to acknowledge Eric Alani and Marcus Smolka for critical reading of the manuscript. We would also like to thank members of the Cornell R3 group and the Crickard laboratory for helpful input during development of the project.

## Author contributions

JH purified proteins, performed experiments, analyzed data, and helped with the writing of the manuscript. BF performed experiments, analyzed data, and helped with writing and editing of the manuscript. JD provided reagents and cloning support. JBC performed experiments, analyzed data, provided reagents, and wrote the manuscript with input from JH and BF. This work was supported by NIGMS R35142457 to JBC and Cornell Startup funds to JBC.

