## Supplemental_Data for "Rad53 regulates the lifetime of Rdh54 at homologous recombination intermediates"

### Supplemental Information
### Supplemental Table 1

| Strains | Genotype | Source or reference |
|---|---|---|
| *LSY-2202-15D (WT)* | *MATa ade2-n his3::NatMX4 met22:klURA3* | *(Mazon et al. 2010)* |
| *LSY-2205-11C (WT)* | *MATalpha ade2-I lys2:GAL-ISCE1 his3:HphMX4* | *(Mazon et al. 2010)* |
| *JBC0011 (15D)* | *RDH54::RDH54-KanMX* | *(Keymakh et al. 2022)* |
| *JBC0013 (11C)* | *RDH54::RDH54-KanMX* | *(Keymakh et al. 2022)* |
| *JBC0014 (15D)* | *RDH54::rdh54K318R* | *(Keymakh et al. 2022)* |
| *JBC0016 (11C)* | *RDH54::rdh54K318R* | *(Keymakh et al. 2022)* |
| *JBC0018 (15D)* | *RDH54::rdh54T851A* | *This study* |
| *JBC0020 (11C)* | *RDH54::rdh54T851A* | *This study* |
| *JBC0026 (15D)* | *RDH54::rdh54S852A* | *This study* |
| *JBC0028 (11C)* | *RDH54::rdh54S852A* | *This study* |
| *JBC0054 (15D)* | *RDH54::rdh54S852D* | *This study* |
| *JBC0060 (11C)* | *RDH54::rdh54S852D* | *This study* |
| *JBC0056 (15D)* | *RDH54::rdh54T851D* | *This study* |
| *JBC0062(11C)* | *RDH54::rdh54T851D* | *This study* |
| *JBC0086 (15D)* | *RDH54::rdh54T851A,S852A* | *This study* |
| *JBC0092 (11C)* | *RDH54::rdh54T851A,S852A* | *This study* |
| *JBC0080 (15D)* | *RDH54::HIS3* | *This study* |
| *JBC0083(11C)* | *RDH54::HIS3* | *This study* |
| *BY4741 (WT)* | | *Dharmacon* |
| *BY4741* | *rdh54Δ* | *Dharmacon* |
| *HK663A-leu2-R1 (A)* | *leu2-EcoRI RAD5+ ade2-1 can1-100 his3-11,15 trp1-1 ura3* | *(Klein H 1997)* |
| *HK6793B-leu2BSTEII (alpha)* | *leu2-BstEII RAD5+ ade2-1 can1-100 his3-11,15 trp1-1 ura3* | *(Klein H 1997)* |
| *JBC206* | *leu2-EcoRI RAD5+ ade2-1 can1-100 his3-11,15 trp1-1::TRP1 ura3* | *This study* |
| *JBC207* | *leu2-BstEII RAD5+ ade2-1 can1-100 his3-11,15 trp1-1 ura3::URA3* | *This study* |
| *JBC0174* | *JBC206 RDH54::RDH54-KanMX* | *This study* |
| *JBC0175* | *JBC206 RDH54::rdh54T851A-KanMX* | *This study* |
| *JBC0176* | *JBC206 RDH54::rdh54T851D-KanMX* | *This study* |
| *JBC0177* | *JBC206 RDH54::rdh54S852A-KanMX* | *This study* |
| *JBC178* | *JBC206 RDH54::S852D-KanMX* | *This study* |

| | | |
|---|---|---|
| *JBC179* | *JBC206 RDH54::rdh54T851A,S852A-KanMX* | *This study* |
| *JBC0180* | *JBC207 RDH54::RDH54-KanMX* | *This study* |
| *JBC0181* | *JBC207 RDH54::rdh54T851A-KanMX* | *This study* |
| *JBC0182* | *JBC207 RDH54::rdh54T851D-KanMX* | *This study* |
| *JBC0183* | *JBC207 RDH54::rdh54S852A-KanMX* | *This study* |
| *JBC0184* | *JBC207 RDH54::S852D-KanMX* | *This study* |
| *JBC0185* | *JBC207 RDH54::rdh54T851A,S852A-KanMX* | *This study* |
| *JBC228* | *JBC206 RDH54::KanMX* | *This study* |
| *JBC229* | *JBC207 RDH54::KanMX* | *This study* |
| *yRA53* | *MAT*a::DEL Hocs::hisG uraD851trp1DEL63 leu2::KAN hmlDEL::hisGHMR::ADE3 ade3::GAL::Hocan1DEL::UR::Hocs::NAT, RA::LEU2, A3::TRP1 | *Anand et al 2014* |
| *yRA163* | *yRA53 RDH54::HPHMX* | *Anand et al 2014* |
| *JBC0084* | *yRA53 leu2::KAN::HPHMX* | *This study* |
| *JBC0099* | *JBC0084 RDH54::RDH54KanMX* | *This study* |
| *JBC0100* | *JBC0084 RDH54::rdh54T851A* | *This study* |
| *JBC0101* | *JBC0084 RDH54::rdh54T851D* | *This study* |
| *JBC0102* | *JBC0084 RDH54::rdh54S852A* | *This study* |
| *JBC0103* | *JBC0084 RDH54::rdh54S852D* | *This study* |
| *JBC0162* | *JBC0084 RDH54::rdh54T851A,S852A* | *This study* |
| *JKM179* | *hoΔ hml::ADE1 MATα hmr::ADE1 ade1-110 leu2,3-112 lys5 trp1::hisG ura3-52 ade3::GAL:HO* | *Kind gift from J. Haber* |
| *JBC0144* | *JKM179 RDH54::RDH54-3xHA-KanMX* | *This study* |
| *JBC0146* | *JKM179 RDH54::rdh54T851A-3xHA-KanMX* | *This study* |
| *JBC0148* | *JKM179 RDH54::rdh54T851D-3xHA-KanMX* | *This study* |
| *JBC0152* | *JKM179 RDH54::rdh54S852A-3xHA-KanMX* | *This study* |
| *JBC0252* | *JKM179 RDH54::rdh54S852D-3xHA-KanMX* | *This study* |
| *JBC0164* | *JKM179 RDH54::rdh54T851A,S852A-3xHA-KanMX* | *This study* |
| *Diploid (JBC0011 X JBC0013)* | *RDH54::RDH54-KanMX/ RDH54::RDH54-KanMX* | *This study* |
| *Diploid (JBC0011XJBC0016)* | *RDH54::RDH54-KanMX /RDH54::rdh54K318R* | *This study* |
| *Diploid (JBC0011XJBC0020)* | *RDH54::RDH54-KanMX/ RDH54::rdh54T851A* | *This study* |
| *Diploid (JBC0018XJBC0020)* | *RDH54::rdh54T851A/ RDH54::rdh54T851A* | *This study* |
| *Diploid (JBC0026X0028)* | *RDH54::rdh54S852A/ RDH54::rdh54S852A* | *This study* |
| *Diploid(JBC0054XJBC0060)* | *RDH54::rdh54S852D/ RDH54::rdh54S852D* | *This study* |
| *Diploid(JBC0056XJBC0062)* | *RDH54::rdh54T851D/RDH54::rdh54T851D* | *This study* |

| | | |
|---|---|---|
| *Diploid (JBC0086XJBC0092)* | *RDH54::rdh54T851A,S852A/RDH54::rdh54T851A,S852A* | *This study* |
| *Diploid (JBC174XJBC180)* | *RDH54::RDH54-KanMX/ RDH54::RDH54-KanMX* | *This study* |
| *Diploid (JBC175XJBC181)* | *RDH54::rdh54T851A/ RDH54::rdh54T851A* | *This study* |
| *Diploid (JBC176XJBC182)* | *RDH54::rdh54T851D/RDH54::rdh54T851D* | *This study* |
| *Diploid (JBC177XJBC183)* | *RDH54::rdh54S852A/ RDH54::rdh54S852A* | *This study* |
| *Diploid (JBC178XJBC184)* | *RDH54::rdh54S852D/ RDH54::rdh54S852D* | *This study* |
| *Diploid (JBC179XJBC185)* | *RDH54::rdh54T851A,S852A/RDH54::rdh54T851A,S852A* | *This study* |
| *Diploid (JBC0080XJBC0083)* | *RDH54::HIS3/RDH54::HIS3* | *This study* |
| *Diploid (JBC228XJBC229)* | *RDH54::KanMX/RDH54::KanMX* | *This study* |

### Supplemental Table 2

| **Plasmid backbone** | **Construct** | **Source** |
|---|---|---|
| pET15 | 6xHis-Rdh54-WT | **This study** |
| pET15 | 6xHis-Rdh54K318R | **This study** |
| pET15 | 6xHis-Rdh54T851A | **This study** |
| pET15 | 6XHis-Rdh54T851D | **This study** |
| pET15 | 6xHis-Rdh54S852A | **This study** |
| pET15 | 6xHis-Rdh54S852D | **This study** |
| pET15 | 6xHis-Rdh54T851AS852A | **This study** |
| pET15 | 6xHis-Rdh54-mCherry-WT | **This study** |
| pET15 | 6xHis-Rdh54-mCherryK318R | **This study** |
| pET15 | 6xHis-Rdh54-mCherryT851A | **This study** |
| pET15 | 6xHis-Rdh54-mCherryS852A | **This study** |
| pET15 | 6xHis-Rdh54-mCherryT851D | **This study** |
| pET15 | 6xHis-Rdh54-mCherryS852D | **This study** |
| pET15 | 6xHis-Rdh54-mCherryT851AS852A | **This study** |
| pRS415 | *RDH54-WT* | **This study** |
| pRS415 | *rdh54S21A* | **This study** |
| pRS415 | *rdh54S167A* | **This study** |
| pRS415 | *rdh54S387A* | **This study** |
| pRS415 | *rdh54S621A* | **This study** |
| pRS415 | *rdh54S790A* | **This study** |
| pRS415 | *rdh54S791A* | **This study** |
| pRS415 | *rdh54T851A* | **This study** |
| pRS415 | *rdh54S852A* | **This study** |
| pRS415 | *rdh54K318R* | **This study** |
| pRS415 | *rdh54K318RT851A* | **This study** |
| pRS305 | *RDH54-KanMX* | **This study** |
| pRS305 | *rdh54T851A-KanMX* | **This study** |
| pRS305 | *rdh54T851D-KanMX* | **This study** |
| pRS305 | *rdh54S852A-KanMX* | **This study** |

| | | |
|---|---|---|
| pRS305 | *rdh54S852D-KanMX* | **This study** |
| pRS305 | *rdh54K318R-KanMX* | **This study** |
| pRS305 | *rdh54T851A,S852A-KanMX* | **This study** |
| pET15 | 6xHis-Rad53 | **A kind gift from Dirk Remus** |
| pET15 | 6xHis-Rad53D339A | **A kind gift from Dirk Remus** |
| pET15 | 6xHis-GFP-Rad53 | **This study** |
| pET15 | 6xHis-GFP-Rad53D339A | **This study** |

### Supplemental Table 3

| Strain | Selective Media | Leu+ Frequency (Total Population) |
|---|---|---|
| WT | 345 | $1.00 \times 10^{-5}$ |
| *rdh54T851A:rdh54T851A* | 42 | $0.45 \times 10^{-5}$ |
| *rdh54T851D:rdh54T851D* | 54 | $1.25 \times 10^{-5}$ |
| *rdh54S852A:rdh54S852A* | 51 | $0.58 \times 10^{-5}$ |
| *rdh54S852D:rdh54S852D* | 243 | $0.77 \times 10^{-5}$ |
| *rdh54T851A,S852A:rdh54T851A,S852A* | 68 | $0.77 \times 10^{-5}$ |
| *rdh54:rdh54* | 258 | $1.00 \times 10^{-5}$ |

### Supplemental Table 4

| Strain | CO | NCO | BIR | Total | Chromosome Loss |
|---|---|---|---|---|---|
| WT | 94 | 72 | 8 | 174 | 0 |
| *rdh54T851A:rdh54T851A* | 76 | 611 | 17 | 704 | 0 |
| *rdh54T851D:rdh54T851D* | 122 | 189 | 32 | 343 | 0 |
| *rdh54S852A:rdh54S852A* | 42 | 93 | 25 | 160 | 0 |
| *rdh54S852D:rdh54S852D* | 74 | 343 | 31 | 448 | 1 |
| *RDH54:rdh54T851A* | 64 | 125 | 7 | 196 | 0 |
| *RDH54:rdh54K318R* | 62 | 125 | 9 | 183 | 0 |
| *rdh54T851A,S852A:rdh54T851A,S852A* | 186 | 181 | 12 | 379 | 0 |

#### Supplemental Table 5

| | | *In vitro* | | | |
|---|---|---|---|---|---|
| *In vivo* SuperPhos | Both | WT-Rad53 | Rad53D339A | Rdh54 No Rad53 (-ATP) | Rdh54 No Rad53 (+ATP) |
| S19 (1) | S19 | S19 | S673 | S158 | S158 |
| S51 | | T23 | | | |
| S165 | | T28 | | | |
| S386 | | T30 | | | |
| S619 | | S48 | | | |
| S790 | | T54 | | | |
| S791 | | S169 | | | |
| T851 (3) | | S189 | | | |
| S852 (57) | | S194 | | | |
| S901 (1) | | T197 | | | |
| | | S456 | | | |
| | | S587 | | | |
| | | T637 | | | |
| | | T691 | | | |
| | | S692 | | | |
| | | S705 | | | |
| | | S738 | | | |
| | | S775 | | | |
| | | S846 | | | |
| | | T847 | | | |
| | | T849 | | | |
| | T851 | T851 | | | |
| | S852 | S852 | | | |
| | | S872 | | | |
| | S901 | S901 | | | |

**Phosphorylated residues identified by mass spectrometry**

#### Supplemental Figure 1

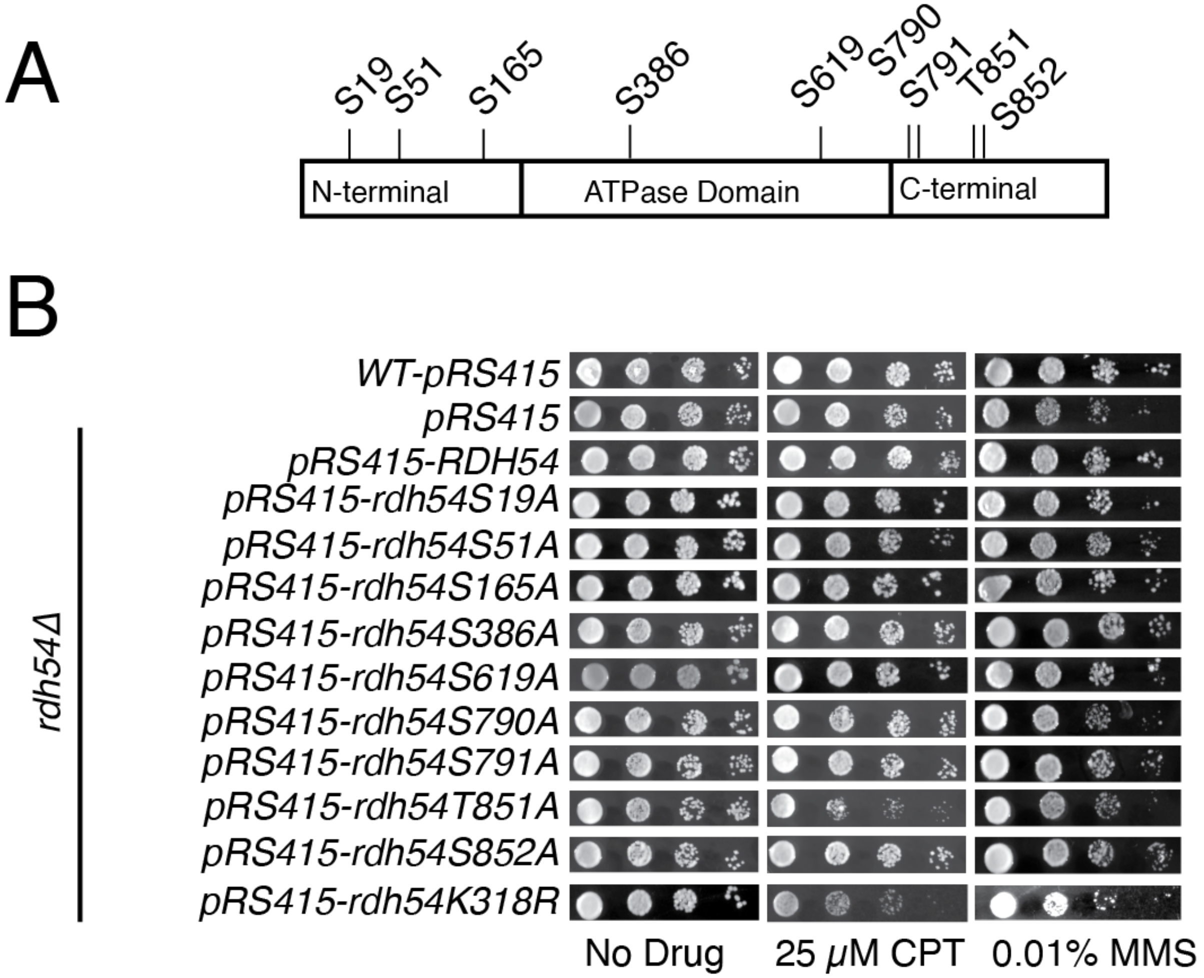


**Supplemental Figure 1: *rdh54T851A* shows enhanced DNA damage phenotype**
**(A).** Schematic diagram illustrating Rdh54 phosphorylation site as identified by the database SuperPhos. All S/T were converted to A. (**B).** Yeast growth assays to measure the growth of different *RDH54* alleles in the presence of 25 µM CPT or 0.01% MMS.

#### Supplemental Figure 2

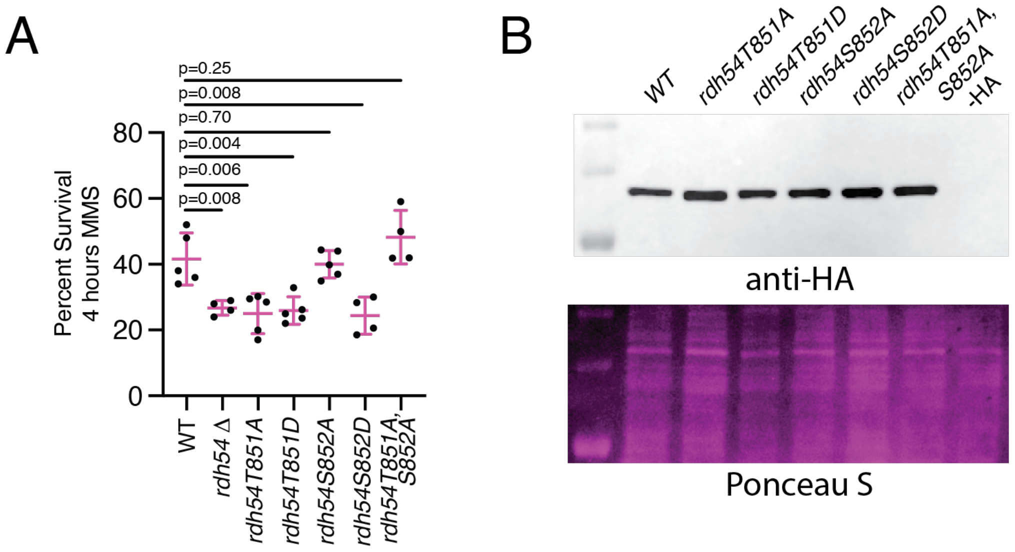


**Supplemental Figure 2: Rdh54 mutants present MMS sensitivity**
**(A).** Graph representing the percent survival of strains treated with 0.01% MMS for 4 hours. The Strains are WT (W303), *rdh54Δ*, *rdh54T851A*, *rdh54T851D*, *rdh54S852A*, *rdh54S852D*, and *rdh54T851A,S852A*. The bars represent the mean of the data, and the error bars represent the standard deviation of at least four independent experiments. P-values were generated using the student's t-test. (B). Western blot of HA tagged versions of RDH54, *rdh54T851A*, *rdh54T851D*, *rdh54S852A*, *rdh54S852D*, and *rdh54T851A,S852A* that illustrates comparable expression.

### Supplemental Figure 3

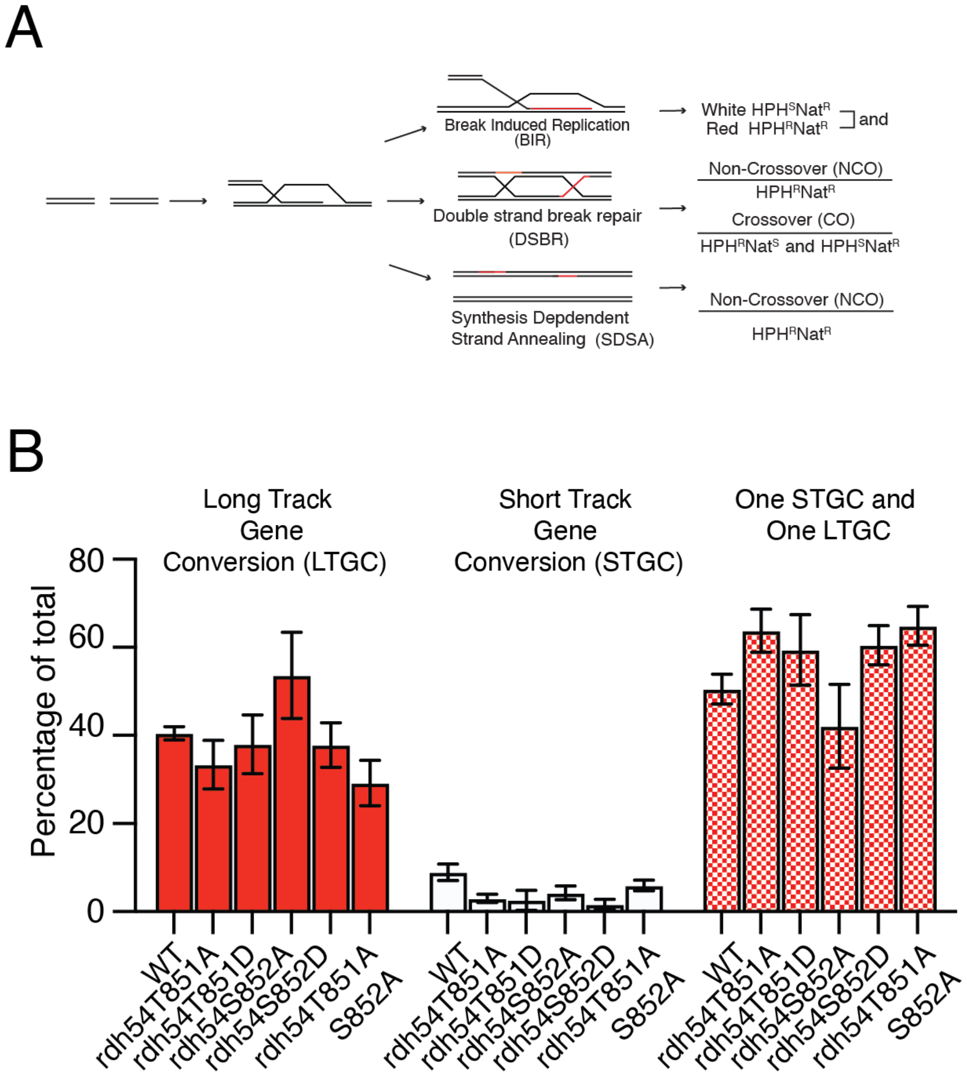


**Supplemental Figure 3: Rdh54 mutants do not affect gene conversion**
**(A).** Schematic outcome depicting the HR outcomes that can be deduced from the red/white assay. These outcomes result in different levels of genetic exchange between chromosomes and are inferred from color and antibiotic sensitivity of specific sectors. **(B).** Bar graph illustrating the populations of solid red colonies (LTGC), solid white colonies (STGC), and mixed sectored colonies (One STGC/ One LTGC). The bars represent the mean, and the error bars represent the standard deviation of at least three independent experiments.

### Supplemental Figure 4

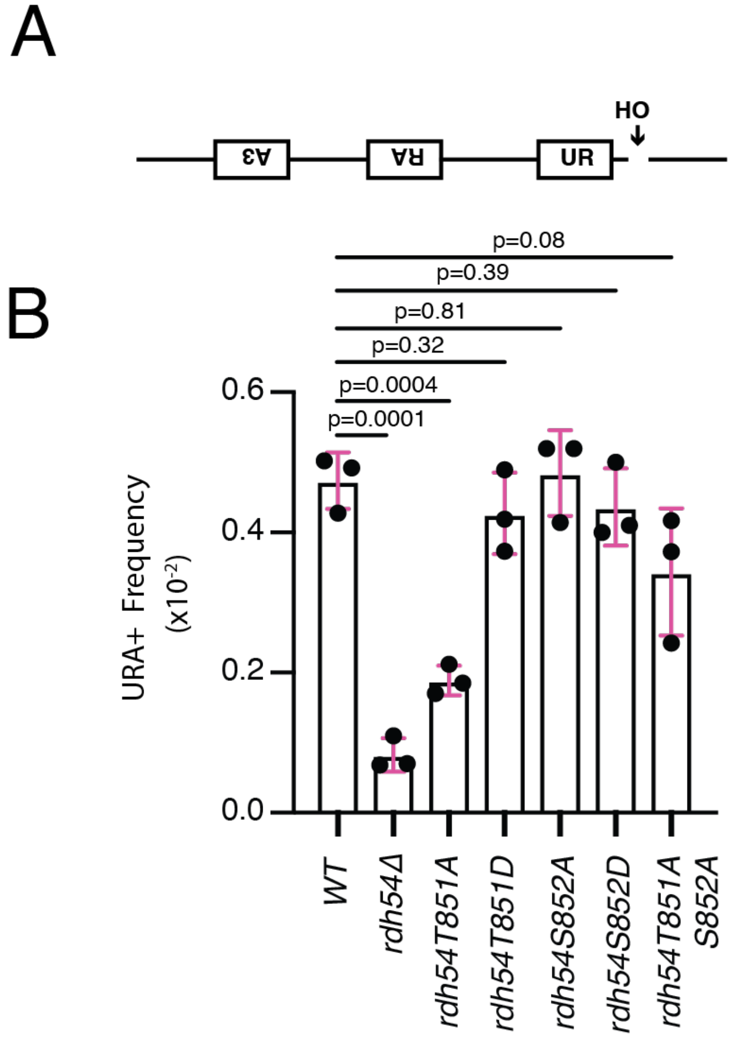


**Supplemental Figure 4: *RDH54* mutants are also defective in DNA template switching**
**(A).** Template defining assay used to measure intrachromosomal template switching during DSBR. The completion of an intact *URA3* gene requires a template switch to occur during repair. **(B).** Measured Ura+ frequency for *RDH54* WT and different alleles of *RDH54*. The bar represents the mean, and the error bars represent the standard deviation for three independent experiments. The p-values were generated by a two tailed t-test.

### Supplemental Figure 5

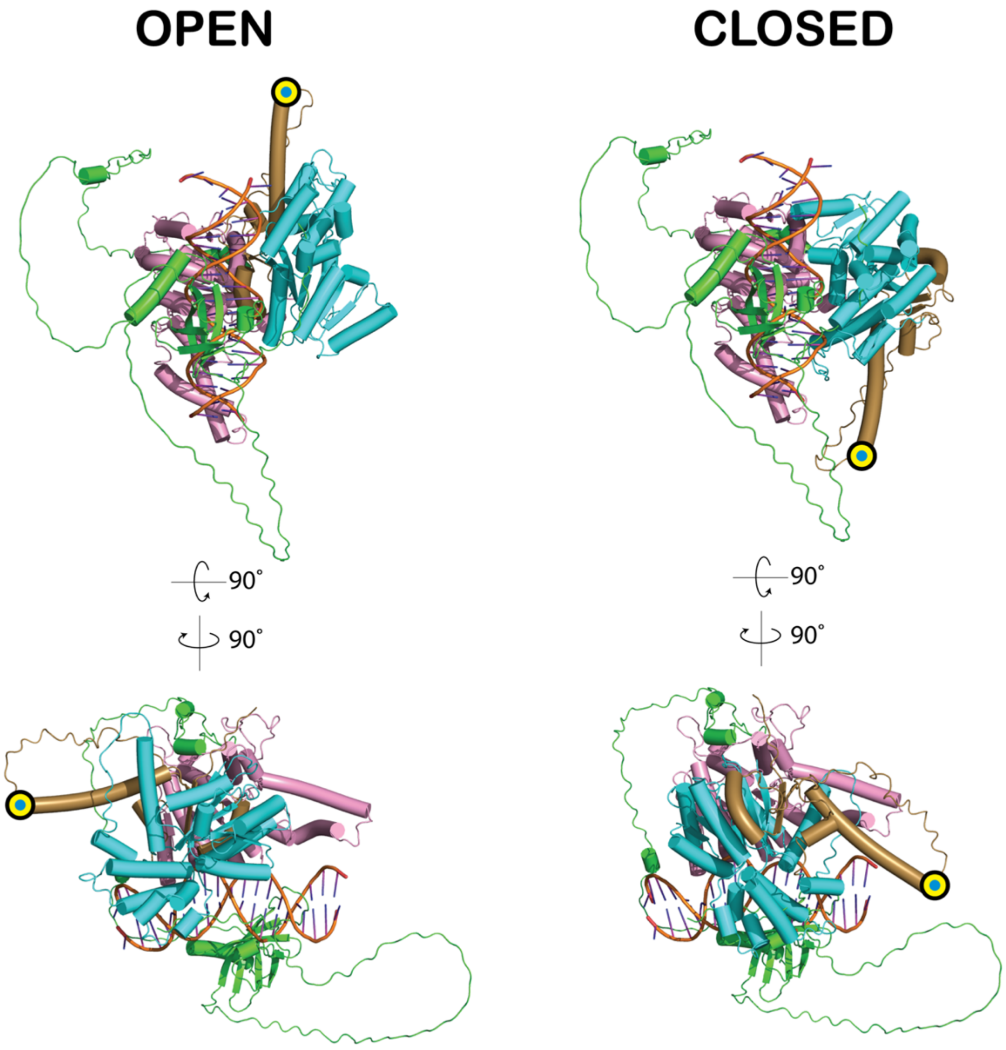


**Supplemental Figure 5: Additional views of the open and closed models of yRdh54**
To generate the open and closed models of yRdh54, the two RecA-like domains in the AlphaFold predicted structure of yRdh54 (https://www.uniprot.org/uniprotkb/B3LN76) were individually superimposed onto the equivalent domains of either the *Sulfalobus solfataricus* SWI2/SNF2 ATPase in the DNA-bound open state (PDB:1Z63), or AMP-PNP:DNA bound PcrA (PDB:3PJR) in the closed state. Most of the N-terminal domain of yRdh54 (green, residues: 1-261) is predicted to be unstructured (per-residue confidence (pLDDT) score < 50) and its position relative to other parts of the model cannot be interpreted. Lobe 1 (residues: 262-259), pink; lobe 2 (residues: 550-807), cyan; C-terminal domain (residues: 808-924), brown. The position of T851 is indicated with a yellow and cyan bullseye. DNA coordinates (orange) are from the *S. solfataricus* ATPase X-ray crystal structure (PDB:1Z63).

### Supplemental Figure 6

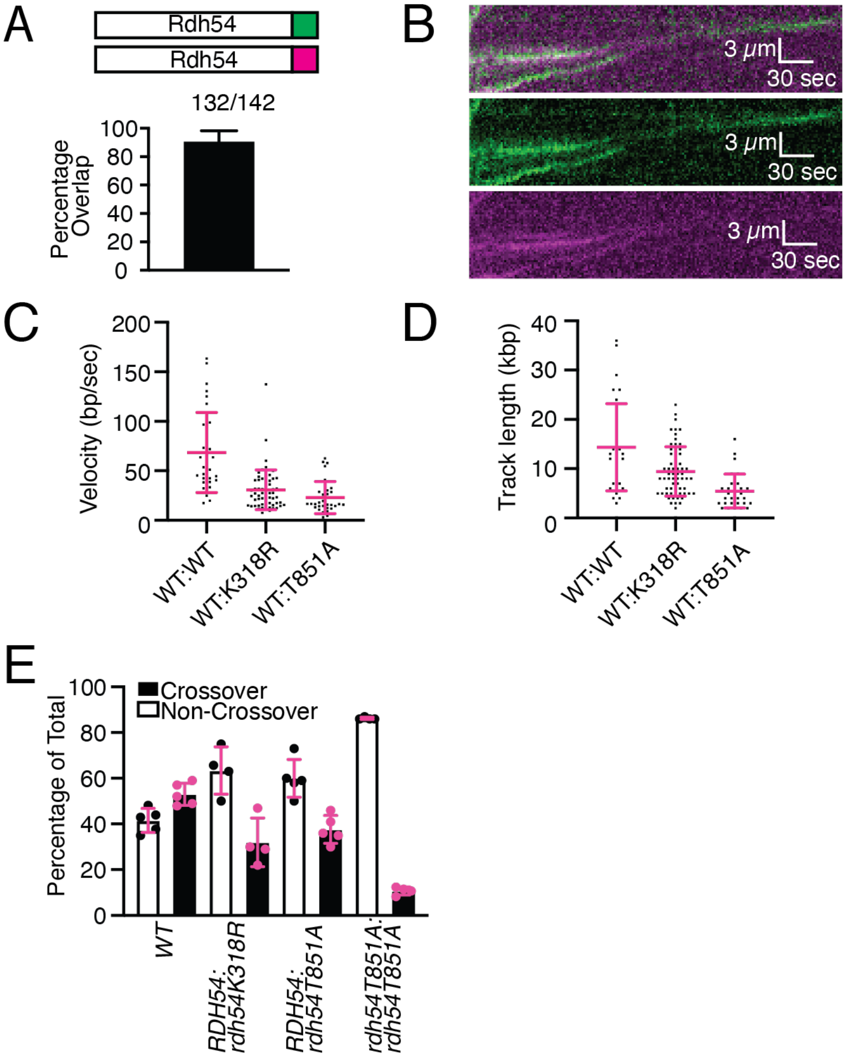


**Supplemental Figure 6: Rdh54 forms a mixed oligomer during translocation**

**(A).** Schematic illustrating experiments performed with equal molar Rdh54-mCherry and Rdh54-GFP (Top). Graphical representation of the number of Rdh54-mCherry molecules that overlap with Rdh54-GFP molecules. The error bar represents the standard deviation of three independent experiments (Bottom). **(B).** Representative kymographs illustrating the co-translocation of Rdh54-mCherry and Rdh54-GFP molecules. The channels are merged (top), magenta (middle), and green (bottom). **(C).** Dot plot representing the velocities (bp/sec) of mixed WT:WT (1:1) (N=32), WT:Rdh54K318R (1:1) (N=62), and WT:Rdh54T851A (1:1) (N=31). The bar and the error bars represent the mean and standard deviation of the data. **(D).** Dot plot of the track lengths (kbp) of mixed WT:WT (1:1) (N=32), WT:Rdh54K318R (1:1) (N=62), and WT:Rdh54T851A (1:1) (N=31). The bar and the error bars represent the mean and standard deviation of the data. **(E).** Percentage of CO and NCO outcomes for WT, *RDH54:rdh54K318R*, and *RDH54:rdh54T851A* strains. The bar represents the mean, and the error bars represent the standard deviation of four independent experiments from four different zygotes.

### Supplemental Figure 7

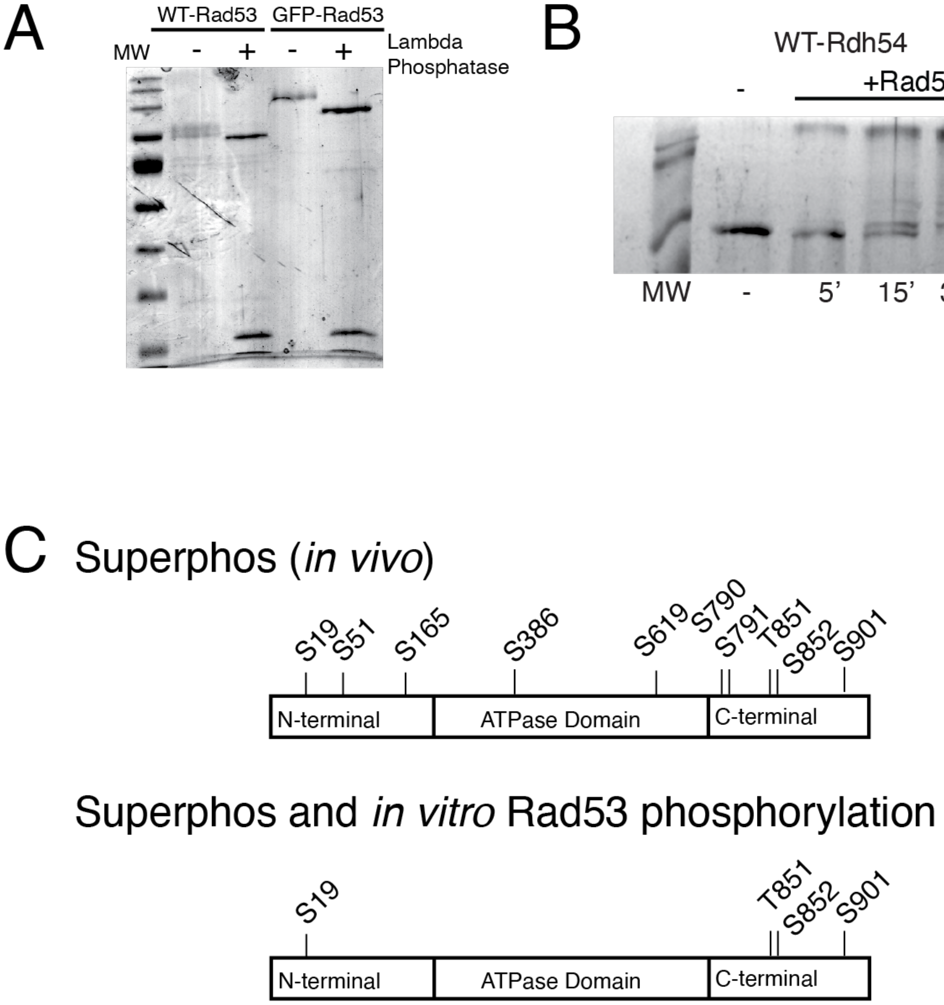


**Supplemental Figure 7: Rdh54 directly interacts with and is a target of Rad53**
**(A).** Representative SDS-PAGE illustrating purified Rad53 and GFP-Rad53 with and without Lambda phosphatase illustrating that Rad53 and GFP-Rad53 are purified in their phosphorylated form. **(B).** SDS-PAGE phostag gel with un-phosphorylated Rdh54, and Rdh54 phosphorylated by Rad53 for 5’, 15’, and 30’. **(C).** Schematic diagram of phosphorylation sites identified in the SuperPhos database (Top, reproduced from Supplemental Figure 1 and Supplemental Table 5) and phosphorylation sites identified both from SuperPhos and from an *in vitro* kinase reaction with Rad53 (Bottom, the data can also be found in Supplemental Table 5).

#### Supplemental Figure 8

A

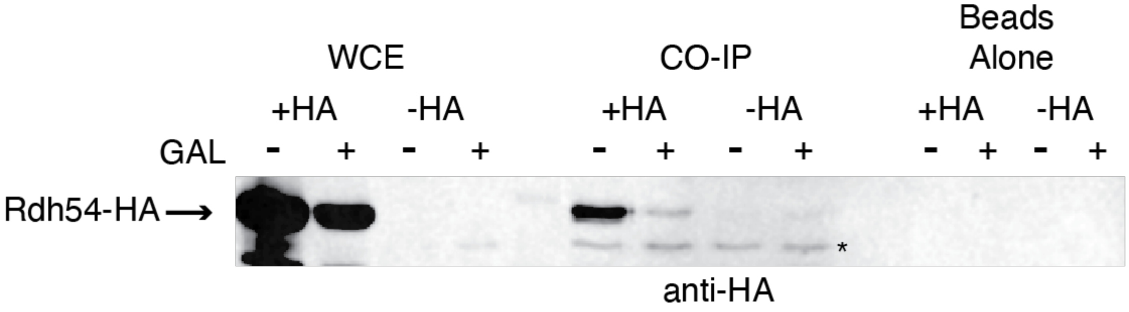


**Supplemental Figure 8: Rdh54 co-immunoprecipitated with Rad53**
**(A).** Representative western blot illustrating the CO-IP of Rdh54-HA with Rad53. Samples shown are from strains with an HA tagged Rdh54 and NO HA tag on the Rdh54. For WCE, IP'd sample and Beads alone. * is an unspecific band.
